# Unveiling the Molecular Architecture of *Candida auris* Ribosome

**DOI:** 10.1101/2025.06.26.661723

**Authors:** A. Atamas, A. Stetsenko, D. Incarnato, A. Rogachev, A. Maciá Valero, S. Billerbeck, A. Guskov

## Abstract

*Candida auris* is an emerging multidrug-resistant fungal pathogen causing life-threatening invasive candidiasis and bloodstream infections (candidemia), posing significant global health challenges. Despite its importance, the protein translation in pathogenic fungi is poorly characterized. Using cryo-electron microscopy and single-particle reconstruction, we resolved high-resolution structures of the 80S ribosome from *C. auris* in its vacant state and in complexes with three inhibitors: cycloheximide (CHX), blasticidin-S (BLS), and geneticin (G418). We uncovered a unique substitution of a key nucleotide in the P-site of the small ribosomal subunit (C1160 in *C. auris*), which may influence ribosome-tRNA interactions and translation fidelity. Comparative analysis of ribosome inhibitor interactions showed that resistance to CHX was observed in only two Candida species examined, while BLS binding displayed no significant differences between *C. auris* and *S. cerevisiae*, although *C. auris* was more sensitive to it. We identified that G418 exhibits promiscuous binding across multiple nonspecific sites, yet its primary interaction site at the decoding center remains highly conserved among Candida species.

These findings provide a previously uncharacterized structure of the *C. auris* ribosome, highlighting novel features that may be leveraged for the development of targeted antifungal therapies to combat multidrug resistance. These insights not only enhance our understanding of ribosomal inhibitor interactions but also suggest potential biomarkers for predicting antifungal susceptibility in clinical applications.

## Introduction

*Candida auris* (current name *Candidozyma auris [1]*) is a species of pathogenic yeast highly resistant to multiple drugs, and it has gained global awareness for its role in causing severe infections and outbreaks within healthcare settings. This fungal pathogen presents a substantial public health concern due to its alarming resistance to medications, rapid transmission, and the potential for exacerbated infections due to various biological and epidemiological factors. Although historically *Candida albicans* has been the predominant species, there has been a noticeable rise in infections caused by non-albicans Candida spp. This increase is thought to be largely driven by the increased usage of antifungal medications, such as fluconazole. Appallingly, infections by non-albicans species tend to exhibit higher mortality rates and demonstrate greater resistance to antifungal drugs compared to *C. albicans* infections. *C. auris*, an emerging and formidable pathogen, has instigated numerous hospital outbreaks worldwide. Its close-to-invincible resistance to drugs severely limits the available treatment options.

*C. auris* was first isolated from the ear discharge of a patient in Japan in 2009 [2]. By 2015, the disease had been registered in 15 countries, and by 2020, this pathogen had been detected in 40 countries [3]. Its emergence and rapid spread accompanied by the development of multidrug resistance in the last decade have marked it as a significant global health threat [4].

One of the possible candidates conferring such extraordinary drug resistance displayed by *C. auris* might be its robust protein translation machinery, and at the moment studies on *C. auris* translational responses to antifungal drugs [5,6] are gaining momentum. Nevertheless, there is a lack of structural / functional insight which proved to be instrumental for the development of antibiotics targeting protein synthesis machinery in numerous pathogens [7–9].

To fill this knowledge gap, we performed a structure / function investigation on translation machinery of *C. auris* in an attempt to identify any peculiarities in the structure of its ribosome, which might be useful not only for the rational drug development but also to evaluate whether it has acquired certain critical evolutionary adaptations. In this study we present single-particle cryo-electron microscopy (Cryo-EM) structures of the vacant *C. auris* ribosome and its complex with three inhibitors cycloheximide (CHX), blasticidin S (BLS) and geneticin (G418) complemented with cell-free translation inhibition assays. Our study revealed significant sensitivity of the ribosome to geneticin and differences in the configuration of ribosomal RNA compared to the *Candida albicans* ribosome.

## Materials and Methods

### Ribosome purification

*C. auris* (ATCC MYA–5001) 80S ribosomes were purified following the protocol in [10].

Candida cells were grown in flasks to OD_600_ (optical density at 600 nm) 1.0 in yeast extract-peptone-dextrose (YPD) medium at 30°C. Cells were pelleted by centrifugation, resuspended with YP, and incubated in flasks with vigorous shaking (250 rpm) for 10.5 min. Cells were pelleted and washed three times in buffer M [30 mM Hepes-KOH (pH 7.5), 50 mM KCl, 10 mM MgCl_2_, 8.5% (w/v) mannitol, 2 mM dithiothreitol (DTT) and 0.5 mM EDTA]. The cell pellet was resuspended in buffer M. The cell suspension was transferred into a 50-ml round-bottom tube (Nalgene) with glass beads (Sigma-Aldrich).

Cells were mechanically disrupted nine times by vortexing at 40 Hz for 1 min with 1 min breaks on ice between each vortex. Glass beads were removed by rapid centrifugation (20,000 g for 2 min), and the lysate was further clarified by longer centrifugation (30,000 g for 9 min). Polyethylene glycol (PEG) 20000 (Hampton Research) was then added to a final concentration of 4.5% (w/v) and the solution was allowed to stand on ice for 5 minutes. The solution was clarified by centrifugation (20,000 g, 5 min). Then the concentration of PEG 20000 was adjusted to 8.5% (w/v) and the solution was left for 10 minutes on ice. The ribosomes were pelleted (17,500 g for 10 min), the supernatant was discarded, and the residual solution was removed by briefly rotating the pellet (14,500 g for 1 min). Ribosomes were suspended (from 8 to 10 mg/ml) in buffer M+ (buffer M with a KCl concentration adjusted to 150 mM, with the addition of protease inhibitors and heparin). Ribosomes were then purified with a 10 to 30% (w/v) sucrose gradient in buffer S [20 mM Hepes-KOH (pH 7.5), 120 mM KCl, 8.3 mM MgCl_2_, 2 mM DTT, and 0.3 mM EDTA] using an SW28 rotor (18000 rpm) at 15 min). Fractions containing the ribosome were assembled. PEG 20000 was then added to a final concentration of 7% (w/v); and the solution was left for 10 min on ice. Ribosomes were precipitated (17,500 g for 10 min), the supernatant was poured off, and the remaining solution was removed by briefly rotating the sediment (17,500 g for 1 min). Ribosomes were suspended (25 mg/ml) in buffer G [10 mM Hepes-KOH (pH 7.5), 50 mM KOAc, 10 mM NH_4_OAc, 2 mM DTT and 5 mM Mg(OAc)_2_].

### rRNA sequencing

For rRNA sequencing, RNA from 10 μg of ribosomes were purified on Monarch RNA Cleanup 10 μg columns (New England Biolabs, cat, T2030L). 1 μg RNA was then mixed with 1 μl 50 μM random hexamers, 2 μl 10 mM dNTPs, and 4 μl of First Strand Buffer 5X [250 mM Tris-HCl, pH 8.3; 375 mM KCl; 15 mM MgCl_2_] in 17 μl final, and incubated at 94°C for 6 minutes to fragment the RNA. Fragmented RNA was then quickly chilled on ice for 2 min, before adding 2 μl 0.1 M DTT, 1 μl TGIRT-III reverse transcriptase (InGex) and 1 μl SUPERaseIn RNase Inhibitor (ThermoFisher Scientific, cat. AM2694). RNA was reverse transcribed for 10 min at 25°C, 30 min at 57°C, and 30 min at 60°C, after which the reverse transcriptase was heat-inactivated by incubating at 75°C for 15 min. RNA was then used as input for the NEBNext® Ultra™ II Non-Directional RNA Second Strand Synthesis Module (New England Biolabs, cat. E6111S), and second strand synthesis was performed as per manufacturer instructions. The resulting dsDNA was then used as input for the NEBNext® Ultra™ II DNA Library Prep Kit for Illumina® kit (New England Biolabs, cat. E7645S) as per manufacturer instructions. The final libraries were sequenced on the Illumina NextSeq 1000 platform. *Sequence analysis*

Protein sequences were retrieved from the UniProt database (https://www.uniprot.org). The following UniProt accession numbers were used:

*S. cerevisiae* - P0CX28, *C. auris* - A0A2H1A644, *C. albicans* - A0A1D8PEV4, *C. parapsilosis* - G8BCU4, *C. glabrata* - Q6FT24, *C. dubliniensis* - B9W952. The sequences and secondary structure diagram rRNA were retrieved from the RNAcentral database (https://rnacentral.org) [11]. The following accession numbers were used:

*S. cerevisiae* 25S - URS000061F377/559292, *C. albicans* 25S - URS00008C81BB/5476, *S. cerevisiae* 18S - URS00005F2C2D/559292, *C. albicans* 18S - URS000059AE3B/237561,

*C. glabrata* 18S - URS0000288C3C/5478, *C. parapsilosis* 18S - URS00021CE91B/5480, *C. dubliniensis* 18S - URS000005D3DF/42374, *H. sapiens* 18S *-* URS0000704D22/9606.

Multiple sequence alignments and matrix were performed using the Clustal Omega tool on the UniProt website, using default parameters [12].

### Cryo-electron microscopy: complex formation, grid freezing and image processing

The purified ribosome sample was filtered (0.22 μm centrifugal filters, Millipore) and concentrated to a final concentration of ∼1–2 mg/mL. Antibiotics were added at the following concentrations: 1 mM BLS and G-418, 0.5 mM CHX. Aliquots of 2.7 μl were applied to carbon grids (Quantifoil Au R1.2/1.3 with ultrafine 2 nm carbon layer, 300 mesh), excess liquid was removed for 3–5 s using an FEI Vitrobot Mark IV (Thermo Fisher Scientific), and samples frozen by immersion in liquid ethane. The prepared grids were transferred to a Titan Krios 300 keV microscope (Thermo Fisher Scientific) equipped with a K3 direct electron detector (Gatan). Zero-loss images were recorded semi-automatically using the UCSF Image script. The GIF-quantum energy filter was adjusted to a slit width of 20 eV. Images were acquired at various nominal magnifications with pixel sizes of 0.836 Å and a defocus range from -0.5 to -2.0 μm. Movies were collected with 50 frames dose-fractionated over 3.5 s.

Motion correction, CTF estimation, manual and template-based particle picking, 2D classification, Ab initio volume generation, CTF global and local refinements, and non-uniform 3D refinement were performed using cryoSPARC (v 4.0)[13]. Using Chimera, the separate masks for the focused refinement were generated for the 60S and 40S subunits. The cryo-EM data processing schemes for the vacant *C. auris* ribosome and in complex with inhibitors are presented in Figs. S5 to S7.

### Modeling

The structure of the *Candida albicans* 80S ribosome [10] was used as a template to build the model. Model-to-map alignment was performed in Chimera [14]. The 60S and 40S subunits were refined separately into their respective focused refined maps using Phenix real-space refinement. Protein and rRNA chains were visually checked in Coot [15] and corrected manually. All studied inhibitors were manually docked into the experimental density, followed by real-space refinement in Phenix [16].

### Inhibition of cell-free translation by CHX, BLS and G-418

We prepared CFTs from *C. auris* strain used for structural analysis. These CFTs were programmed with spLUC mRNA and then their capacity to produce the enzyme in the presence of increasing concentrations of inhibitors was examined. *C. auris* CTF preparation and translation reactions were assayed for LUC activity as described in the method 1 protocol in [17]. Capped and polyadenylated spLUC mRNA was prepared by T7 transcription of EcoRI-linearized plasmid pQQ101 encoding spLUC [18]. *C. auris* extract programmed with 10 ng spLUC RNA were incubated at 25°C in the absence or presence of various concentrations of inhibitors for 100 min and spLUC activity was then measured luminometrically. Cell-free transition experiments were carried out in triplicates. IC_50_ values are presented the text as mean ± SD. Plots of normalized activity were presented as mean ± SEM. GraphPad Prism 10 software was used to perform all statistical tests and to make plots.

### Figure preparation

Panels of figures showing structural models were prepared using Chimera [19]. The sequence of logo drawings was made using Adobe Illustrator.

## Results and discussion

The 18S ribosomal RNA gene, part of the small ribosomal subunit, is one of the most widely used markers in phylogenetic studies due to its conserved nature across eukaryotes, while also containing variable regions that enable species differentiation [20–22].

At first, we performed bioinformatics analysis of rRNA in different Candida species, however, even the curated Candida genome database (http://www.candidagenome.org) lacks the complete sequence of rRNA from *C. auris*.

To overcome this, we performed in-house sequencing of *C. auris* rRNAs. A preliminary reference was built by scanning the *C. auris* genome with the eukaryotic small and large subunit rRNA covariance models from RFAM [23] to identify the 18S and 25S sequences. Reads from rRNA sequencing were then mapped onto this reference, and the identified mutations were annotated to define the actual rRNA sequences. The obtained sequences can be found in Supplementary Information file.

Since Candida is a complicated taxonomical genus, as it is polyphyletic and contains yeasts belonging to different families/lineages, it is somewhat less straightforward to analyze the conservation among different Candida spp. [24]. Still, we decided to make such a comparison for those organisms, which are clearly labeled as pathogenic (*C. glabrata (*current name *Nakaseomyces glabratus* [25]*), C. parapsilosis, C. dubliniensis, C. albicans* and *C. auris (*current name *Candidozyma auris* [1])). Within this subgroup, Candida species show a high level of identity with the wild type yeast Saccharomyces, amounting to at least 95% based on the 18S RNA identity matrix. And among each other, their mutual similarity also reaches at least 95%. Intriguingly, the lowest level of identity is observed in *C. auris* and amounts to no more than 92.68% (Table 1).

**Table 1.** Matrix of percent identity of 18S ribosomal RNA.

|  | *S. cerevisiae* | *C. glabrata* | *C. parapsilosis* | *C. dubliniensis* | *C. albicans* | *C. auris* |
| --- | --- | --- | --- | --- | --- | --- |
| *S. cerevisiae* | 100% | 98.44% | 96.29% | 95.04% | 95.79% | **91.27%** |
| *C. glabrata* | 98.44% | 100% | 95.89% | 95.04% | 95.79% | **91.95%** |
| *C. parapsilosis* | 96.29% | 95.89% | 100% | 98.48% | 99.16% | **92.39%** |
| *C. dubliniensis* | 95.04% | 95.04% | 98.48% | 100% | 98.93% | **91.69%** |
| *C. albicans* | 95.79% | 95.79% | 99.16% | 98.93% | 100% | **92.68%** |
| *C. auris* | **91.27%** | **91.95%** | **92.39%** | **91.69%** | **92.68%** | **100%** |

Such differences in identity are normally due to (or within) expansion segments (ES), in which sequence variations are often observed [26]. However, when comparing *C. auris* with *C. albicans* and *S. cerevisiae*, sequence variation can be observed not only in the expansion regions but also in certain rRNA loops (Fig. S1). Changes are observed not only in the 18S rRNA regions, but also in the 25S rRNA sequences (Fig. S2). In some areas, quite significant differences in segment construction, such as shortening or lengthening, are observed. Several examples are described in detail in the supplementary materials (Supplementary material S3 and S4).

Extension segments are believed to confer an improved stability, regulation and expand functionality – there are studies that have shown their effect on ribosome assembly [27] and translation [26,28]. Given the fact that ESs often have multiple contacts with ribosomal proteins [29,30], it is perhaps not very surprising that they can indeed have an impact on the stability of the ribosome structure and the elongation process.

In stark contrast to variability in ESs and in certain rRNA loops, other elements of rRNA, especially those involved in ribosome assembly and function, remain highly conserved.

However, despite the significant conservation of rRNA regions involved in the formation of functional sites, *C. auris* exhibits a change in a certain key nucleotide in the P-site (Fig. 1). In particular, the nucleotide U1191, present in the 18S rRNA of *S. cerevisiae*, other yeasts, and also in humans, located within the h31 helix, is one of the key elements of the ribosomal RNA [31]. The pseudouridylation of this nucleotide facilitates the gradual folding of the tertiary structure of the small subunit’s head in eukaryotes, which forms the P-site [32,33].

**Fig. 1.**
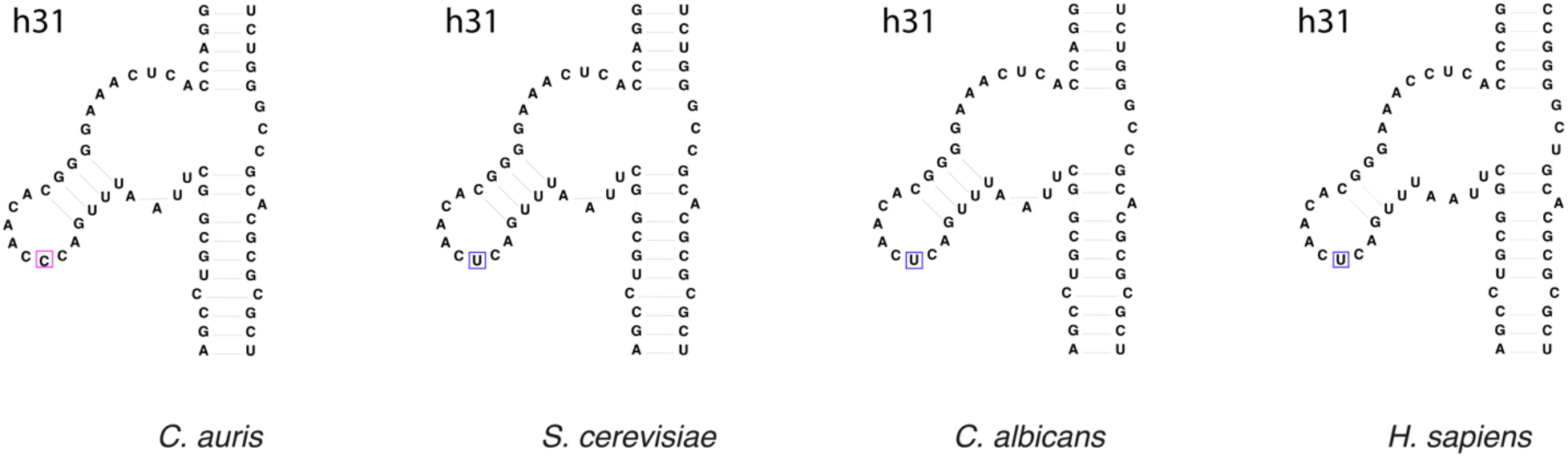
Schematic representation of helix h31 in *C. auris, S. cerevisiae, C. albicans* and *H. sapiens*. The color indicates the nucleotides involved in the P-site of the small ribosomal subunit. In *C. auris*, the key nucleotide in the P-site of the small ribosomal unit is cytosine (pink box), while in the other organisms it is uracil (blue box).

Given its functional role, this nucleotide is usually conserved; however, in the case of *C. auris*, this nucleotide is replaced by cytosine (at least in 84% of cases according to our sequencing), which could potentially affect not only the folding of the small ribosome subunit but also the stability of its head. The presence of cytosine at this particular position is atypical even for bacteria, which typically have guanine at this site [34,35], and replacement of this guanine (G966) with cytosine in *E. coli* results in a slight decrease in ribosome activity [36].

Tetracyclines inhibit tRNA binding to the ribosome and typically have multiple binding sites, with the primary site being the A-site of the small ribosomal subunit [37]. Antibiotics such as tetracycline and tigecycline interact with phosphate groups in helices h31 and h34 via magnesium ions. Thus, these inhibitors do not need specific nucleotides for interaction, making them effective antibiotics with a broad spectrum of action [38]. Nevertheless, it has been shown by Gerrits et al. that replacing nucleotides AGA926-928 in *Helicobacter pylori* (UGC965-967 in *E. coli*, CCC1159-1162 in *C. auris*) with UUC leads to tetracycline resistance [39]. Whether the cytosine triplet in *C. auris* affects tetracycline efficacy or ribosomal activity remains to be elucidated and represents a challenging but potentially informative direction for future research.

To better understand the characteristics of the *C. auris* ribosome, we decided to conduct structural studies of its interaction with different inhibitors. For this analysis, we selected three inhibitors, each of which interacts with different functional sites on the ribosome.

One of the best-known inhibitors of eukaryotic translation is cycloheximide (CHX); it blocks the elongation phase by binding to the E-site [40]. The P56Q substitution in the eL42 protein, as in *C. albicans* [10], prevents binding of CHX to the 60S subunit through the steric hindrance between glutamine and the glutamide group of the inhibitor. This mutation is absent in *C. auris* (at least in the strain we used in our analysis), similarly to *S. cerevisiae* and other Candida species except *C. dubliniensis* (Fig. 2A, 2B). Furthermore, E-site in *C. auris* ribosome is invariantly conserved, hence its sensitivity to CHX should be comparable to that of *S. cerevisiae*. Moreover, *C. auris* has no changes in the rRNA helices surrounding the E-site or in ribosomal proteins in contact with it that could potentially interfere with binding to CHX. This is indeed the case as confirmed by both the cell-free assays and the structure obtained (Fig. 2C, 2D). CFTs in the extracts of *S. cerevisiae* and *C. auris* were reduced by 50% at about 1 μM and completely inhibited at concentration of ∼ 2.5 μM.

**Fig. 2.**
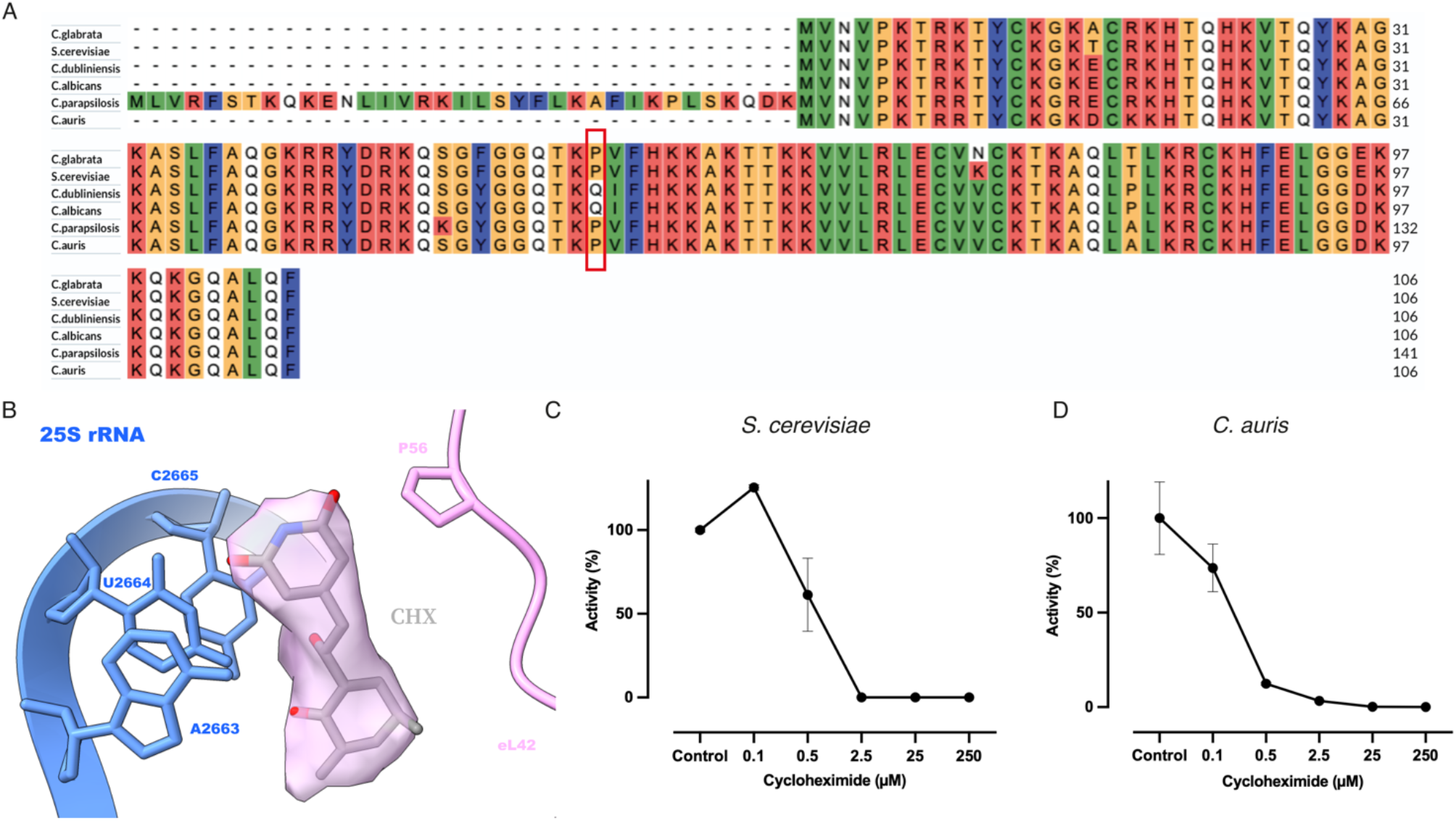
Binding of CHX to the *C. auris* ribosome. (A) Sequence of ribosomal protein L42. *C. albicans* and *C. dubliniensis* exhibit glutamine at position 56 (red box), which protects the E-site nucleotides from being blocked by cycloheximide. The other species contain proline, which does not interfere with the interaction of the inhibitor with the ribosome. (B) Cryo-EM density for cycloheximide (CHX) binding to the *C. auris* ribosome (∼2 Å). (C) Cell-free translation in *S. cerevisiae* extract with cycloheximide. IC_50_: 0.62±0.27 µM. (D) Cell-free translation in *C. auris* extract with cycloheximide. IC_50_: 0.17±0.07 µM.

To elucidate the underlying reasons for this sensitivity of *C. auris* to CHX, we conducted a comparative analysis with the CHX binding site in the *S. cerevisiae* ribosome [PDB: 7N8B]. In the *C. auris* ribosome, the inhibitor binds to the nitrogenous bases of nucleotides U2664 and C2665, the phosphate group of nucleotide C94, and the ribose of nucleotide G93 (Fig. 3A). In *S. cerevisiae*, CHX forms the same contacts (G92, C93, U2763 and C2764 in *S. cerevisiae*) (Fig. 3B); however, in *C. auris*, the inhibitor additionally establishes a contact with the spermidine molecule (SPD) and interacts with a water molecule. Moreover, four out of five hydrogen bonds with nucleoitides in *C. auris* are shorter by at least 0.3 Å. Superimposition of cycloheximide (CHX) molecules revealed that in *C. auris*, the CHX molecule adopts a slightly different conformation, penetrating approximately 1 Å deeper into the ribosome than in *S. cerevisiae* (Fig. 3C, 3D), which perhaps can explain a slightly tighter binding of CHX to *C. auris* ribosome in comparison to *S. cerevisiae* ribosome.

**Fig. 3.**
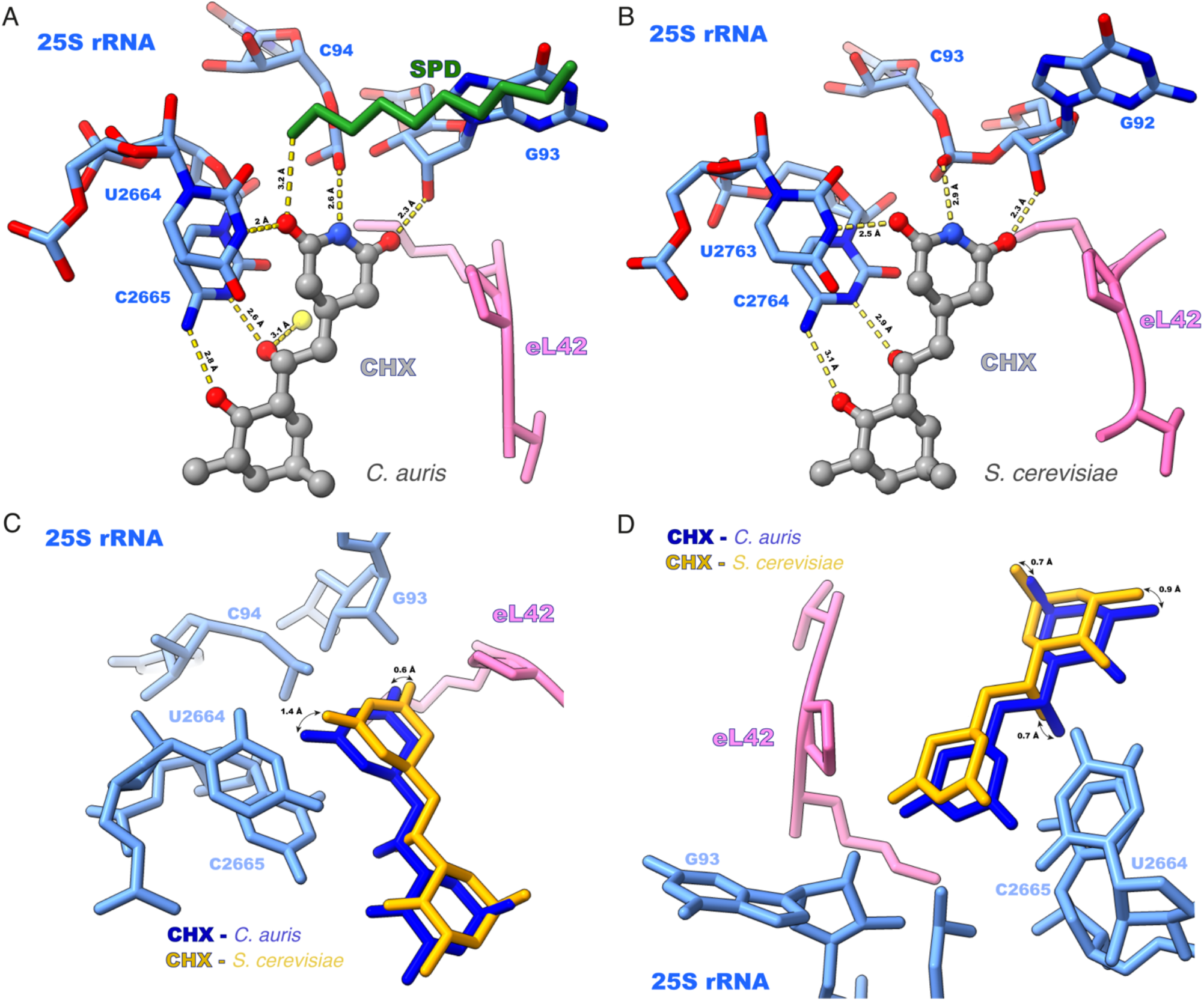
Binding of CHX to the *C. auris* and *S. cerevisiae* ribosome. Close-up views of the *C. auris* (A) and *S. cerevisiae* (B) CHX binding sites. Hydrogen bonds shown as yellow dashed lines. (C and D) Comparison of cycloheximide conformations in *C. auris* (blue) and *S. cerevisiae* (yellow) ribosomes in two orientations.

Blasticidin S (BLS) acts as an inhibitor that modulates P-site activity by interfering with the peptidyltransferase reaction [41]. In both prokaryotes and eukaryotes, its primary interaction is with the guanine of the H82 loop, which replaces the interaction with the 3′-CCA tRNA [42,43].

Despite the complete identity of the P-site between *C. auris, S. cerevisiae* and Candida sp., our cell-free translation assays revealed that *C. auris* ribosomes are significantly more sensitive to blasticidin. Specifically, translation in *C. auris* lysates was fully inhibited at 250 µM of blasticidin (Fig. 4A), while *S. cerevisiae* translation remained partially active (∼40%) at the same concentration (Fig. 4B).

**Fig. 4.**
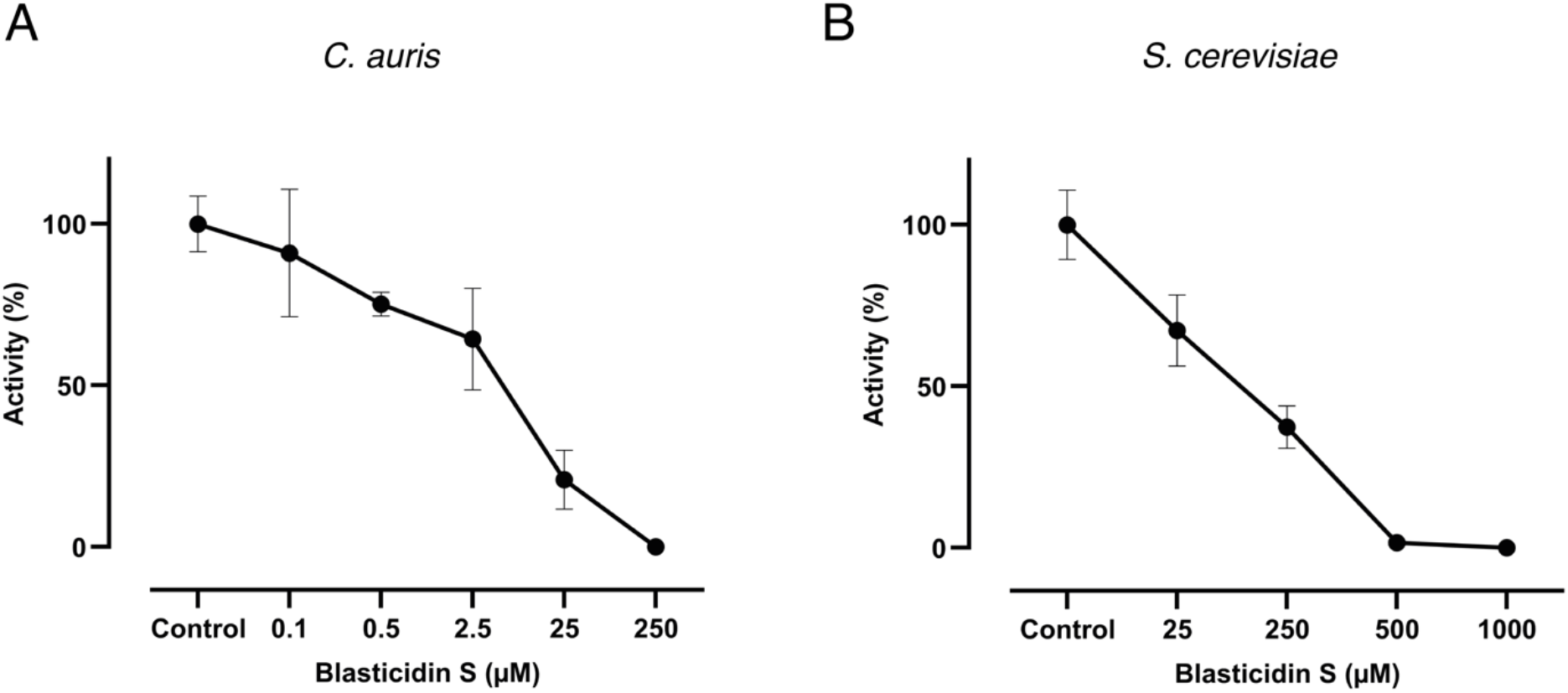
Cell-free translation with the addition of blasticidin S. (A) in *C. auris*. IC_50_: 5.3±3.3 µM (B) in *S. cerevisiae*. IC_50_: 154±94 µM

In an attempt to understand the reasons for this sensitivity, we analyzed the structures of the ribosome with BLS. Initially we aimed to obtain the complex of *C. auris* ribosome with three antibiotics simultaneously: CHX, BLS and G418; however, no density could be assigned to a single BLS molecule. We argued that perhaps conformational changes induced by CHX and G418 disfavored BLS binding, hence we solved the structure of *C. auris* ribosome only in complex with BLS.

This structure revealed minor differences in blasticidin binding to *C. auris* ribosome. In *C. auris*, binding occurs to the nitrogenous base of nucleotide G2520, the phosphate residues of nucleotides A2709 and A2870, and the sugar residue of nucleotide C2321 (Fig. 5A). In the ribosome of *S. cerevisiae* [PDB 4U56] BLS binds to the nitrogenous base of nucleotide G2619 and the phosphate residues of nucleotides C244, C2406, A2808, and A2969 (Fig. 5B).

**Fig. 5.**
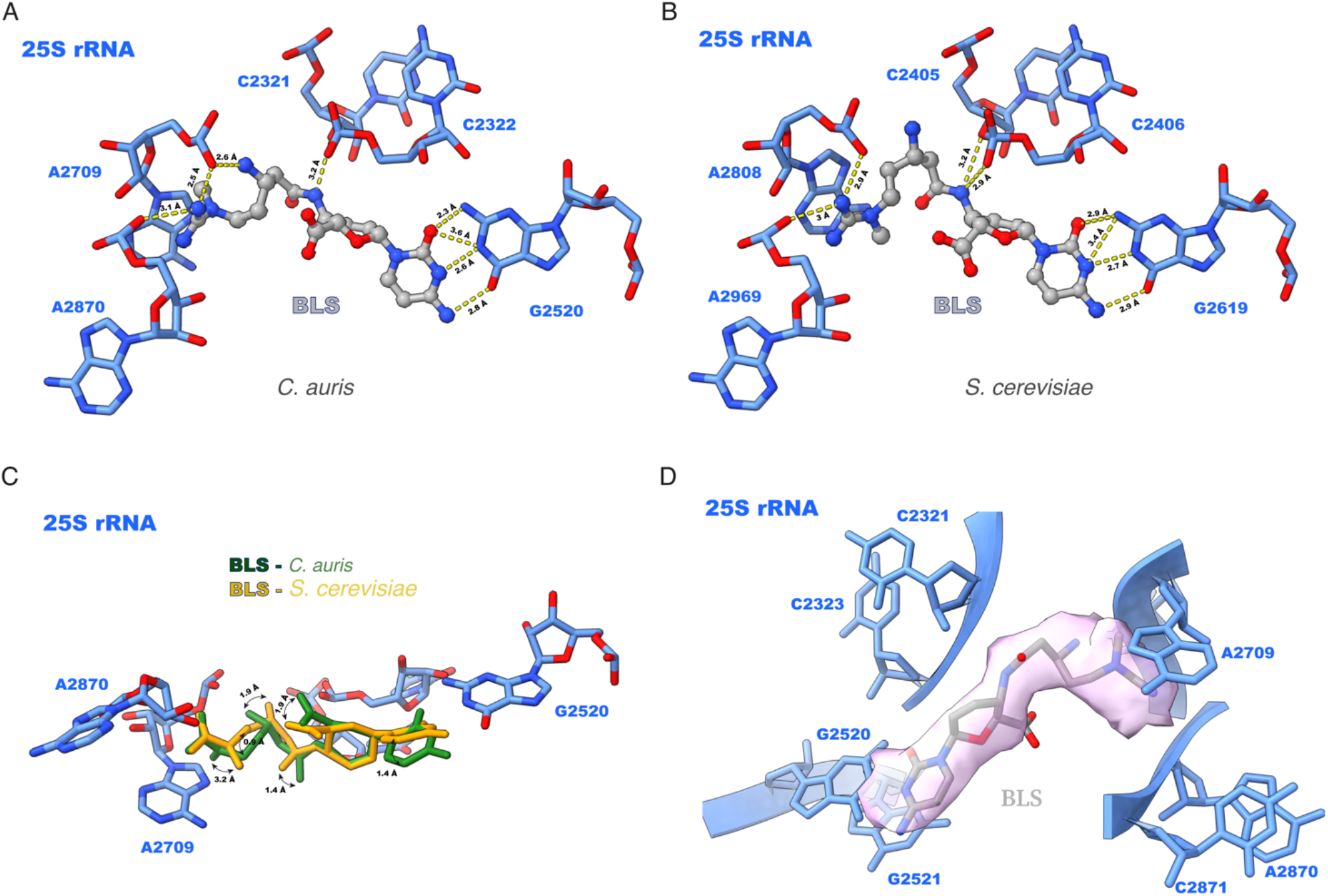
Binding of BLS to the *C. auris* and *S. cerevisiae* ribosomes. Close-up views of the *C. auris* (A) and *S. cerevisiae* (B) BLS binding sites. Hydrogen bonds shown as yellow dashed lines. (C) Comparison of blasticidin conformations in *C. auris* (green) and *S. cerevisiae* (orange) ribosomes. (D) Cryo-EM density for blasticidin S (BLS) binding to the *C. auris* ribosome (∼3 Å).

Superimposing the structures reveals differences in the conformation of the inhibitor’s tail, notably deviation of atoms C5, C6, C9, C10 and C13 (Fig. 5C). However, the density for this region is less well resolved to precisely determine the positions of these atoms (Fig. 5D), hence it cannot be ruled out that this deviation is a modeling artefact. Anyhow, such differences in the BLS interactions with rRNA observed between *S. cerevisiae* and *C. auris* cannot explain the difference in inhibition. Notably, the relatively weak density observed for BLS in *C. auris* may indicate transient or unstable binding, raising the possibility that this is not the sole binding site-although no additional density consistent with BLS was detected elsewhere in the structure.

Aminoglycosides (AG) are a group of broad-spectrum antibiotics that includes streptomycin, neomycin, geneticin (G418) and others [44]. AG have multiple binding sites, thereby exerting diverse effects on the translation mechanism, however they show a high affinity for the A-site of the small subunit of the ribosome, binding to which prevents tRNA translocation [45–47].

A study by Prokhorova et al., [48] on the interaction of aminoglycosides with *S. cerevisiae* ribosome showed that G418 has several binding sites such as: E-site, A-site and the peptide tunnel. Thus, it can affect not only tRNAs but also interfere with the pathway of nascent peptides. Interaction with the A-site of the ribosome is achieved by binding of geneticin to helix h44 in the small subunit. The h44 helix sits in the decoding center of the ribosome, which is responsible for the accuracy of codon-anticodon interactions. The binding of G418 to h44 causes a change in the position of some conserved nucleotides (A1755 and A1756 in *S. cerevisiae*), shifting them in the opposite direction.

In the current study, the *C. auris* ribosome was found to be highly sensitive to G418 compared to *S. cerevisiae*. As can be seen from cell-free translation assays with *C. auris* extract, translation was already reduced by 50% at 3 nM, and at 0.5 μM translation was completely inhibited (Fig. 6A); while *S. cerevisiae* translation was inhibited 50% only at 0.46 μM (Fig. 6B).

**Fig. 6.**
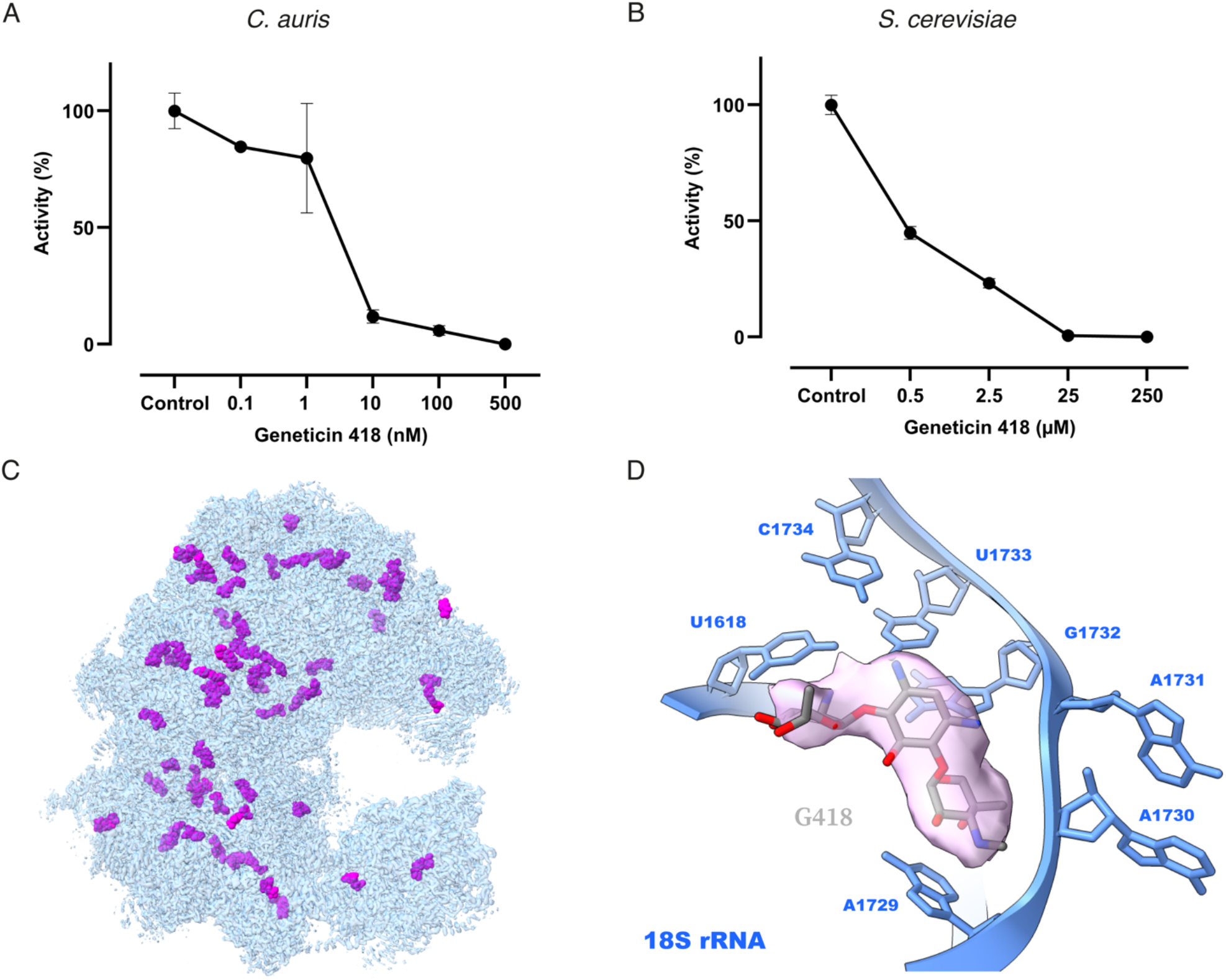
Binding of G418 to the *C. auris* ribosome. (A) Cell-free translation in *C. auris* extracts with geneticin, IC_50_: 2.9±1.5 nM. (B) Cell-free translation in *S. cerevisiae* extracts with geneticin, IC_50_: 0.46±0.07 µM. (C) Overview of the G418 binding sites (purple) in the *C. auris* 80S ribosome. (D) Cryo-EM density for G418 binding to the *C. auris* ribosome in decoding center h44 (∼3 Å resolution).

We identified more than 55 possible binding sites of G418 in our structure (Fig. 6C; Table S1). Geneticin binds to near the peptide tunnel regions and exhibits some interactions with the E- and A-sites, and binding at and near the decoding center. It also binds to the 5.8S rRNA and 5S rRNA segments and interacts with ribosomal proteins including L3, L10, L19, L26, L33 S2, S8 and S13. Upon analyzing the geneticin binding sites involving ribosomal proteins, no significant differences were observed in the interacting amino acids among the pathogenic Candida species examined (*C. glabrata, C. parapsilosis, C. dubliniensis, C. albicans* and *C. auris*). This suggests that these regions interact with the inhibitor in a similar manner across different Candida species. The primary geneticin binding site, namely the decoding center, also remains unchanged (Fig. 6D). However, our results show that G418 has numerous binding sites on the ribosome, which may ensure low nM IC_50_ value, but the reasons for this heightened sensitivity remain to be investigated in future studies.

*C. auris* is one of the few fungal pathogens that is listed by WHO as a pathogen requiring immediate attention due to its alarming resistance to multiple antifungal drugs and rapid spread in healthcare settings. Since 2019, it has been classified as an urgent threat by the CDC in the United States, and since 2022, C. auris, along with *C. albicans*, has been considered a critical priority on the first list of fungi that pose a health threat [49].

These features of *C. auris* highlight the urgent need for new antifungal strategies. Given that for many bacterial pathogens a ribosome is proven to serve as a very successful target, we ventured into structural and functional characterization of ribosomes from fungal pathogens – as even minor differences from human ribosomes can be potentially exploited for the development of a broad-spectrum antifungal antibiotic.

Through the obtained sequencing data, it was possible to identify that ribosomal RNA 18S of *C. auris* has the lowest percentage of identity with another pathogenic Candida species. This is quite unusual because the essential role of 18S rRNA in the translation process is linked to the fact that its sequence changes extremely slowly on evolutionary time scales [50]. Although the percent identity is still above 90%, which is the typical identity within a single genus, we observe multiple changes in the secondary structure of the rRNA. These changes affect not only the expansion segments, where multiple sequence variations usually occur, but also some helices, and not only in 18S rRNA but also in 25S rRNA.

The most unexpected finding is a change in one of the key nucleotides of the P-site in the small subunit of the ribosome. Nucleotide U1191 (in *S. cerevisiae*) in helix h31 is a functional P-site nucleotide and controls the folding of the small subunit head, but unlike other yeasts, in *C. auris* this nucleotide is replaced by cytosine (C1160). It is not yet possible to postulate how this replacement affects the performance of the P-site, but this substitution could potentially influence the interaction dynamics between the ribosome and the tRNA, as well as the overall fidelity and efficiency of translation. Experiments specifically targeting this region of the small subunit are needed to fully understand the impact of this nucleotide.

Based on our data with cycloheximide (CHX), blasticidin-S (BLS) and geneticin (G418), we conclude that only two Candida species are resistant to cycloheximide, while other pathogenic species can be inhibited by blocking the E-site. We found minor differences in BLS binding between *C. auris* and *S. cerevisia*e, with the inhibitor interacting similarly at key ribosomal sites. Despite these similarities, *C. auris* appears to be more sensitive, likely due to interactions with other ribosomal regions that we were not yet able to detect. Furthermore, geneticin binding analysis showed multiple binding sites revealing its promiscuity and the primary geneticin binding site, the decoding center, remains unchanged, suggesting a conserved mode of interaction.

Our experiments showed that the *C. auris* ribosome is sensitive enough to certain types of inhibitors that they can be used as building blocks for developing new antifungal therapies, whereas ribosomal RNA sequencing data will help make these therapies targeted because of the distinct rRNA regions compared to other pathogens and humans.

## Supporting information

Supplemental rRNA sequence

## Acknowledgments

We thank M. Yusupov and Y. Zgadzay for their valuable advice on ribosome preparation; I. Sorokin and Matthew S. Sachs for their advice on working with the cell-free translation system; M. Fraaije and V. Szymanski for an access to Multimode Reader BioTek Synergy H1; and the personnel of NeCEN for their assistance with the data collection.

## Notes

### Competing Interest Statement

The authors have declared no competing interest.

## Reference

1. Liu F, Hu Z-D, Zhao X-M et al. Phylogenomic analysis of the Candida auris-Candida haemuli clade and related taxa in the Metschnikowiaceae, and proposal of thirteen new genera, fifty-five new combinations and nine new species. Persoonia -Mol Phylogeny Evol Fungi 2024;52:22–43.

2. Satoh K, Makimura K, Hasumi Y et al. Candida auris sp. nov., a novel ascomycetous yeast isolated from the external ear canal of an inpatient in a Japanese hospital. Microbiol Immunol 2009;53:41–4.

3. Du H, Bing J, Hu T et al. Candida auris: Epidemiology, biology, antifungal resistance, and virulence. PLOS Pathog 2020;16:e1008921.

4. Kordalewska M, Perlin DS. Identification of Drug Resistant Candida auris. Front Microbiol 2019;10:1918.

5. Zamith-Miranda D, Amatuzzi RF, Munhoz da Rocha IF et al. Transcriptional and translational landscape of Candida auris in response to caspofungin. Comput Struct Biotechnol J 2021;19:5264–77.

6. Iyer KR, Whitesell L, Porco Jr. JA et al. Translation Inhibition by Rocaglates Activates a Species-Specific Cell Death Program in the Emerging Fungal Pathogen Candida auris. mBio 202AD;11, DOI: 10.1128/mbio.03329-19.

7. Jones RN, Fritsche TR, Sader HS et al. Activity of Retapamulin (SB-275833), a Novel Pleuromutilin, against Selected Resistant Gram-Positive Cocci. Antimicrob Agents Chemother 2006;50:2583–6.

8. Serio AW, Keepers T, Andrews L et al. Aminoglycoside Revival: Review of a Historically Important Class of Antimicrobials Undergoing Rejuvenation. EcoSal Plus 2018;8:10.1128/ecosalplus.ESP-0002-2018.

9. Nguyen M, Chung EP. Telithromycin: The first ketolide antimicrobial. Clin Ther 2005;27:1144–63.

10. Zgadzay Y, Kolosova O, Stetsenko A et al. E-site drug specificity of the human pathogen Candida albicans ribosome. Sci Adv 2022;8.

11. The RNAcentral Consortium, Sweeney BA, Petrov AI et al. RNAcentral: a hub of information for non-coding RNA sequences. Nucleic Acids Res 2019;47:D221–9.

12. The UniProt Consortium, Bateman A, Martin M-J et al. UniProt: the Universal Protein Knowledgebase in 2025. Nucleic Acids Res 2025;53:D609–17.

13. Punjani A, Rubinstein JL, Fleet DJ et al. cryoSPARC: algorithms for rapid unsupervised cryo-EM structure determination. Nat Methods 2017;14:290–6.

14. Pettersen EF, Goddard TD, Huang CC et al. UCSF Chimera—A visualization system for exploratory research and analysis. J Comput Chem 2004;25:1605–12.

15. Casañal A, Lohkamp B, Emsley P. Current developments in Coot for macromolecular model building of Electron Cryo-microscopy and Crystallographic Data. Protein Sci 2019;29:1055–64.

16. Liebschner D, Afonine PV, Baker ML et al. Macromolecular structure determination using X-rays, neutrons and electrons: recent developments in Phenix. Acta Crystallogr Sect Struct Biol 2019;75:861–77.

17. Wu C, Amrani N, Jacobson A et al. The Use of Fungal In Vitro Systems for Studying Translational Regulation. In: Lorsch J (ed.). Methods in Enzymology. Vol 429. Academic Press, 2007, 203–25.

18. Wang Z, Fang, Peng, and Sachs MS. The Evolutionarily Conserved Eukaryotic Arginine Attenuator Peptide Regulates the Movement of Ribosomes That Have Translated It. Mol Cell Biol 1998;18:7528–36.

19. Meng EC, Goddard TD, Pettersen EF et al. UCSF CHIMERAX : Tools for structure building and analysis. Protein Sci 2023;32:e4792.

20. Meyer A, Todt C, Mikkelsen NT et al. Fast evolving 18S rRNA sequences from Solenogastres (Mollusca) resist standard PCR amplification and give new insights into mollusk substitution rate heterogeneity. BMC Evol Biol 2010;10:70.

21. Zheng X, He Z, Wang C et al. Evaluation of different primers of the 18S rRNA gene to profile amoeba communities in environmental samples. Water Biol Secur 2022;1:100057.

22. Jamy M, Foster R, Barbera P et al. Long-read metabarcoding of the eukaryotic rDNA operon to phylogenetically and taxonomically resolve environmental diversity. Mol Ecol Resour 2020;20:429–43.

23. Kalvari I, Nawrocki EP, Ontiveros-Palacios N et al. Rfam 14: expanded coverage of metagenomic, viral and microRNA families. Nucleic Acids Res 2021;49:D192–200.

24. Gabaldón T, Naranjo-Ortíz MA, Marcet-Houben M. Evolutionary genomics of yeast pathogens in the Saccharomycotina. FEMS Yeast Res 2016;16.

25. Takashima M, Sugita T. Taxonomy of Pathogenic Yeasts Candida, Cryptococcus, Malassezia, and Trichosporon. Med Mycol J 2022;63:119–32.

26. Hariharan N, Ghosh S, Palakodeti D. The story of rRNA expansion segments: Finding functionality amidst diversity. WIREs RNA 2023;14:e1732.

27. Ramesh M, Woolford JL. Eukaryote-specific rRNA expansion segments function in ribosome biogenesis. RNA 2016;22:1153–62.

28. Ohmayer U, Gil-Hernández Á, Sauert M et al. Studies on the Coordination of Ribosomal Protein Assembly Events Involved in Processing and Stabilization of Yeast Early Large Ribosomal Subunit Precursors. PLOS ONE 2015;10:e0143768.

29. Rhodin MHJ, Dinman JD. A flexible loop in yeast ribosomal protein L11 coordinates P-site tRNA binding. Nucleic Acids Res 2010;38:8377–89.

30. Kisly I, Gulay SP, Mäeorg U et al. The Functional Role of eL19 and eB12 Intersubunit Bridge in the Eukaryotic Ribosome. J Mol Biol 2016;428:2203–16.

31. Xie Q, Wang Y, Lin J et al. Potential Key Bases of Ribosomal RNA to Kingdom-Specific Spectra of Antibiotic Susceptibility and the Possible Archaeal Origin of Eukaryotes. PLOS ONE 2012;7:e29468.

32. Huang H, Karbstein K. Assembly factors chaperone ribosomal RNA folding by isolating helical junctions that are prone to misfolding. Proc Natl Acad Sci 2021;118:e2101164118.

33. Meyer B, Wurm JP, Kötter P et al. The Bowen–Conradi syndrome protein Nep1 (Emg1) has a dual role in eukaryotic ribosome biogenesis, as an essential assembly factor and in the methylation of Ψ1191 in yeast 18S rRNA. Nucleic Acids Res 2011;39:1526–37.

34. Jemiolo DK, Taurence JS, Giese S. Mutations in 16S rRNA in Escherichia coli at methyl-modified sites: G966, C967, and G1207. Nucleic Acids Res 1991;19:4259–65.

35. Jiang J, Seo H, Chow CS. Post-transcriptional Modifications Modulate rRNA Structure and Ligand Interactions. Acc Chem Res 2016;49:893–901.

36. Saraiya AA, Lamichhane TN, Chow CS et al. Identification and role of functionally important motifs in the 970 loop of Escherichia coli 16S ribosomal RNA. J Mol Biol 2008;376:645–57.

37. Pioletti M, Schlünzen F, Harms J et al. Crystal structures of complexes of the small ribosomal subunit with tetracycline, edeine and IF3. EMBO J 2001;20:1829–39.

38. Wilson DN. Ribosome-targeting antibiotics and mechanisms of bacterial resistance. Nat Rev Microbiol 2014;12:35–48.

39. Gerrits MM, de Zoete MR, Arents NLA et al. 16S rRNA Mutation-Mediated Tetracycline Resistance in Helicobacter pylori. Antimicrob Agents Chemother 2002;46:2996–3000.

40. Schneider-Poetsch T, Ju J, Eyler DE et al. Inhibition of eukaryotic translation elongation by cycloheximide and lactimidomycin. Nat Chem Biol 2010;6:209–17.

41. Powers KT, Stevenson-Jones F, Yadav SKN et al. Blasticidin S inhibits mammalian translation and enhances production of protein encoded by nonsense mRNA. Nucleic Acids Res 2021;49:7665–79.

42. Nelli MR, Heitmeier KN, Looper RE. Dissecting the Nucleoside Antibiotics as Universal Translation Inhibitors. Acc Chem Res 2021;54:2798–811.

43. Garreau de Loubresse N, Prokhorova I, Holtkamp W et al. Structural basis for the inhibition of the eukaryotic ribosome. Nature 2014;513:517–22.

44. Mingeot-Leclercq M-P, Glupczynski Y, Tulkens P. Aminoglycosides: Activity and Resistance. Antimicrob Agents Chemother 1999;43:727–37.

45. Garneau-Tsodikova S, J. Labby K. Mechanisms of resistance to aminoglycoside antibiotics: overview and perspectives. MedChemComm 2016;7:11–27.

46. Krause KM, Serio AW, Kane TR et al. Aminoglycosides: An Overview. Cold Spring Harb Perspect Med 2016;6:a027029.

47. Dibrov SM, Parsons J, Hermann T. A model for the study of ligand binding to the ribosomal RNA helix h44. Nucleic Acids Res 2010;38:4458–65.

48. Prokhorova I, Altman RB, Djumagulov M et al. Aminoglycoside interactions and impacts on the eukaryotic ribosome. Proc Natl Acad Sci 2017;114:E10899–908.

49. World Health Organization. WHO Fungal Priority Pathogens List to Guide Research, Development and Public Health Action. World Health Organization, 2022.

50. Pánek J, Kolář M, Vohradský J et al. An evolutionary conserved pattern of 18S rRNA sequence complementarity to mRNA 5′ UTRs and its implications for eukaryotic gene translation regulation. Nucleic Acids Res 2013;41:7625–34.

