## Supplemental data for "Unveiling the Molecular Architecture of *Candida auris* Ribosome"

**Supplementary material**

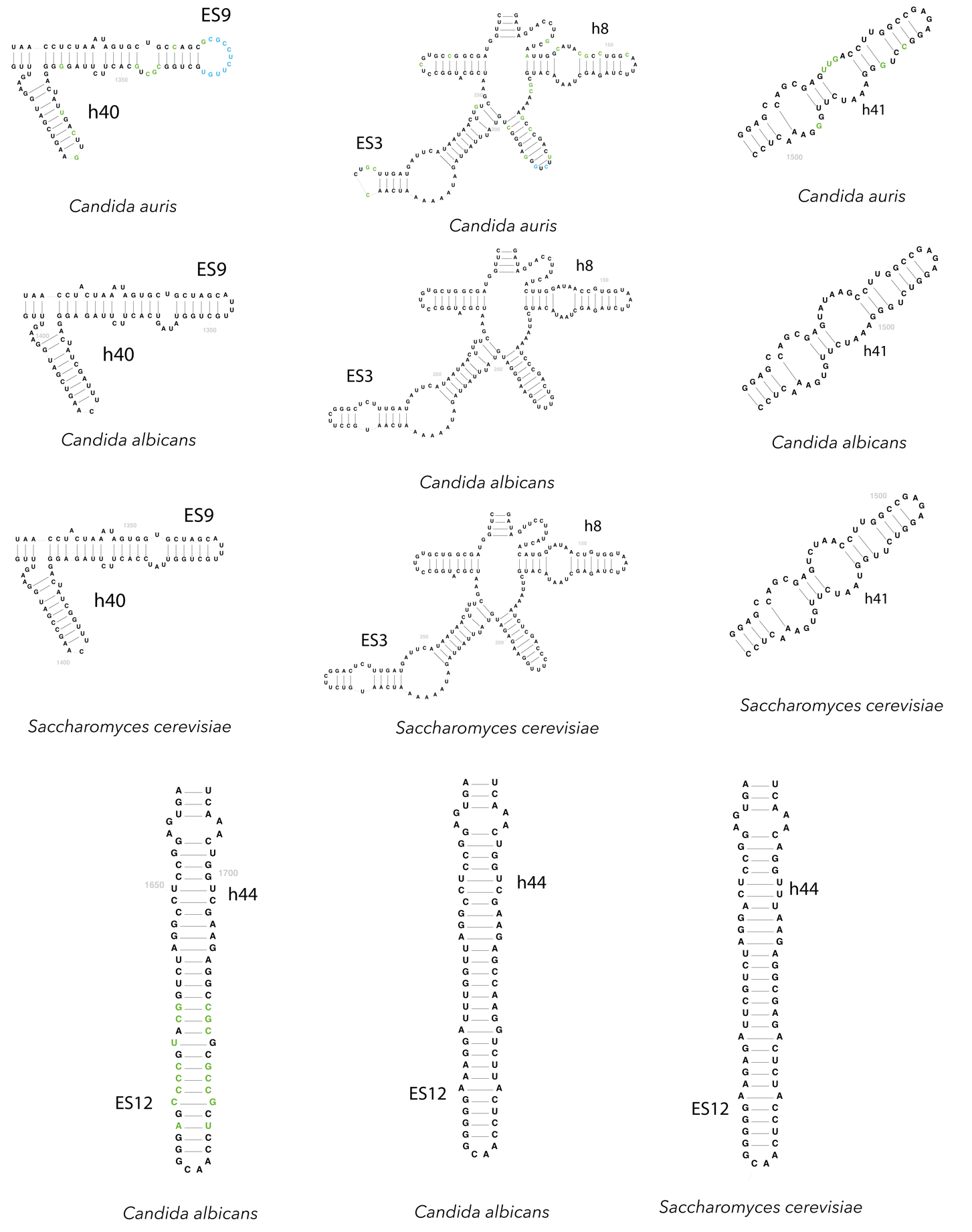

Fig. S1. Comparison of 18S rRNA segments in *C. auris*, *C. albicans* and *S. cerevisiae.* The color indicates a *C. auris* nucleotide that differs from the *C. albicans* and *S. cerevisiae* nucleotide: green - altered nucleotide; blue - additional nucleotide.

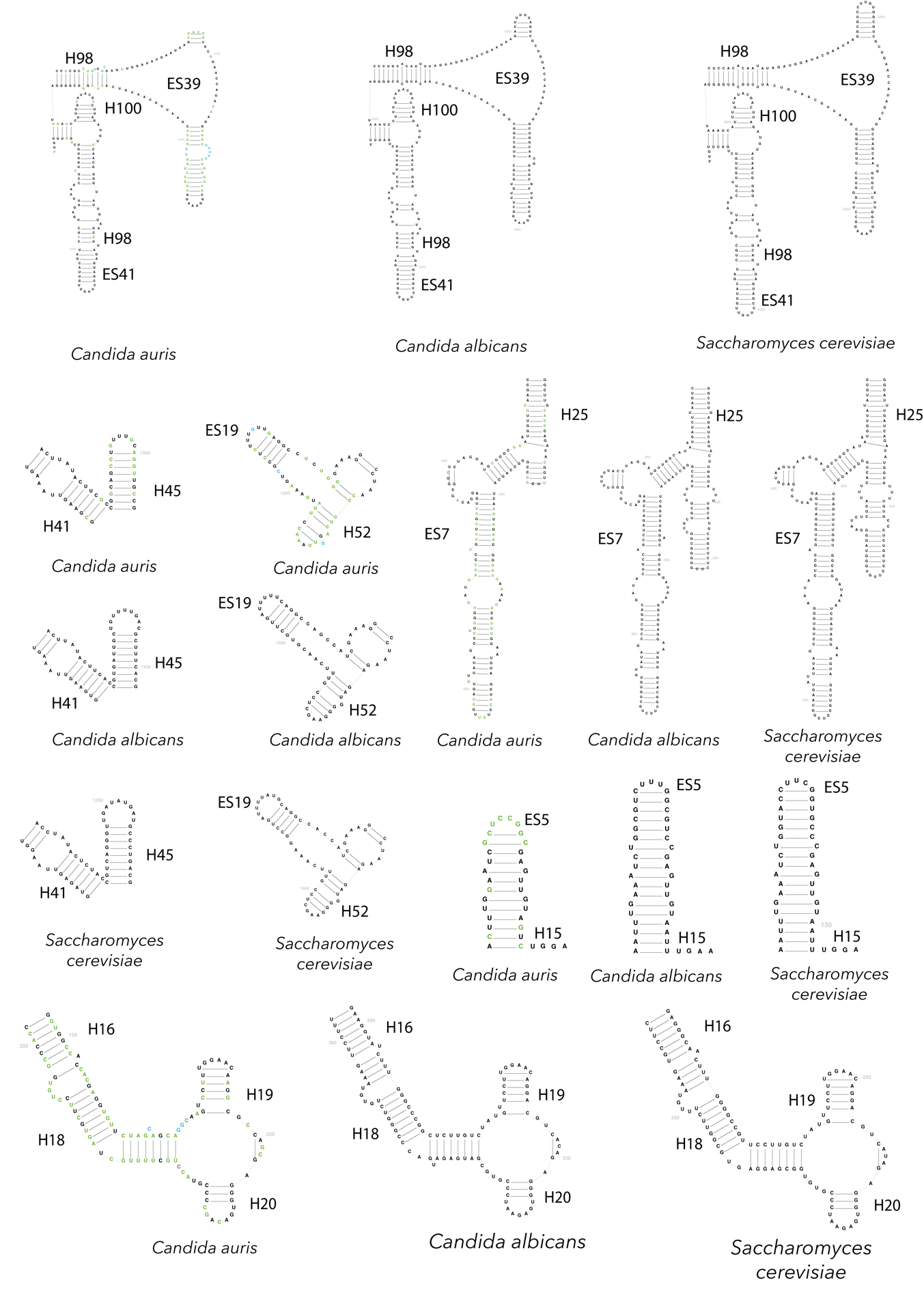

Fig. S2. Comparison of 25S rRNA segments in *C. auris*, *C. albicans* and *S. cerevisiae.* The color indicates a *C. auris* nucleotide that differs from the *C. albicans* and *S. cerevisiae* nucleotide: green - altered nucleotide; blue - additional nucleotide.

**Supplementary material S3**

The ES6 segment performs several important functions, since it plays a role in mRNA recognition[1,2], and also forms a platform for binding elongation factors through interaction with the ES3 helix[3,4] (Fig. S3A). In yeast, this platform is constructed from the interaction of eight nucleotides from the ES6 segment (GUUGGUUU) and eight nucleotides from the ES3 segment (AAAUCAAU) (Fig.S3B). In *C. auris*, the ES3 segment is shortened, and the interaction occurs at the end of the helix rather than near the end as in other yeasts. The last nucleotide of the sequence to which the ES6 segment binds is altered (AAAUCAAC). Since these interactions are also involved in the formation of inter-subunit bridges, it can be hypothesized that such a shift in the interaction site towards the end of the helix might affect the ratcheting movement[5].

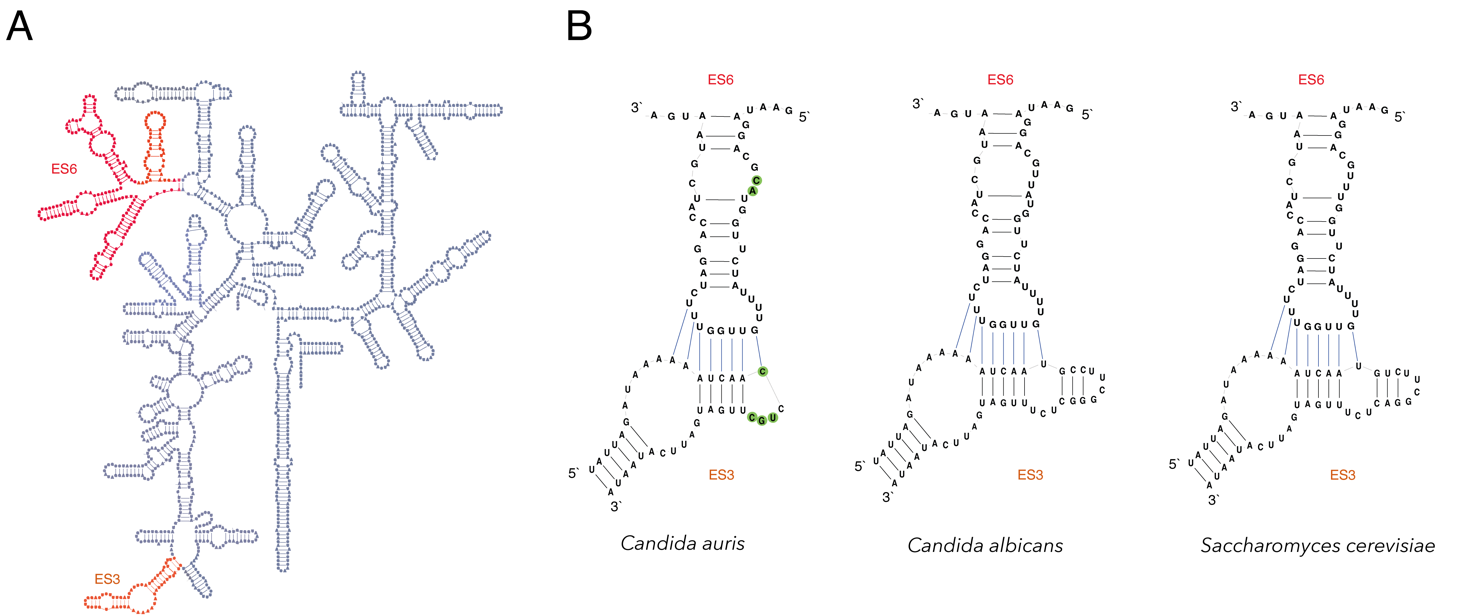

Fig. S3. Schematic representation of the ES3 (orange) and ES6 (red) extension segments. (A) Secondary structure diagram of the 18S rRNA of *S. cerevisiae* (from the RNAcentral database).

(B) Schematic representation of the interaction of the ES3 and ES6 expansion segments in *C. auris*, *C. albicans* and *S. cerevisiae*. The green color indicates the nucleotide of *C. auris*, which is different from that of the nucleotide in *C. albicans* and *S. cerevisiae*.

**Supplementary material S4**

Interestingly Gulay et al., showed that ES27 and ES7 segments have a potential role in the control of elongation[6]. Thus, ES7, interacting with H42-H44, plays a role in binding to elongation factors, and conformational rearrangements of ES27 contribute to the ribosome dynamics, as a result of which the small subunit makes a turn during elongation.

Comparison of these segments with *S. cerevisiae* and *С.* *albicans*, revealed that the tip of the ES27 helix in *С. auris* is ten nucleotides longer than in С. *albicans* and six nucleotides shorter than in *S. cerevisiae* (Fig. S4A)*.* In a study by Fujii et al. it was shown that the ES27 segment serves as a platform for ribosome binding to the enzyme methionine aminopeptidase and shortening of helix length in *S. cerevisiae* affects decoding accuracy, which results in mild tolerance to cycloheximide and anisomycin[7].

Furthermore, the ES7 "shoulder" (site G539-U557 in *C. auris*, G628-U652 in *C. albicans*, G589-U612 in *S. cerevisiae*) is shorter by at least five nucleotides in *C. auris* (Fig. S4B). Similar shortening of the shoulder has been observed in the thermophilic filamentous fungus *Chaetomium thermophilum*, however the impact of such shortening has not been discussed[8]. It is also noteworthy that near this arm there is a cleavage site for endonucleases that are activated by oxidative stress[9]. The ES7 segment has already been proposed as a drug target[10]. Researchers have proposed peptidic aminosugar conjugates that specifically bind to this segment in *C. albicans* and have low affinity for the human ribosome. The study showed that these conjugates have multiple binding sites on the ES7 segment, including the shoulder region. Thus, this segment may be of interest for further studies of the pathogenicity of *C. auris*.

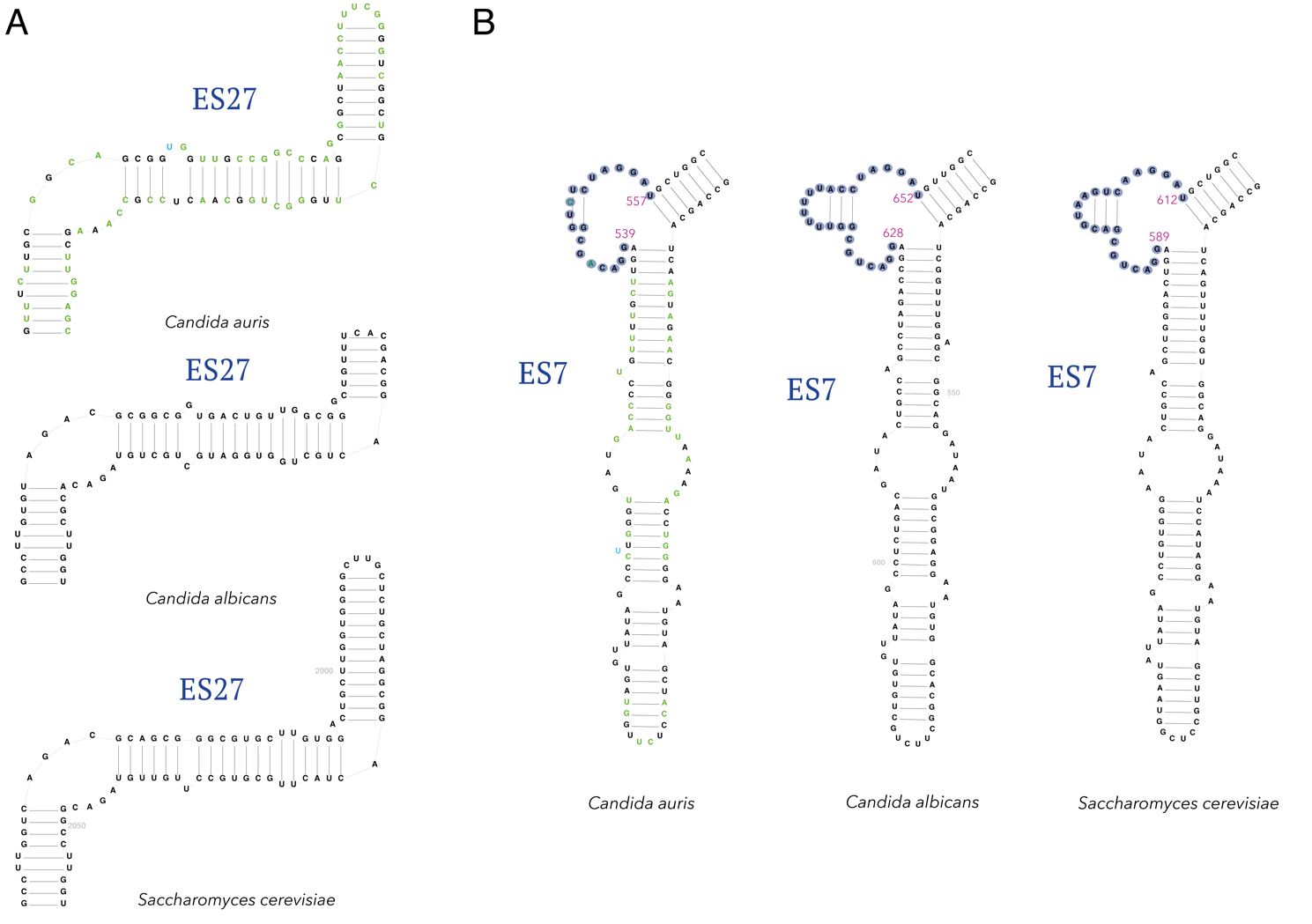

Fig. S4. Schematic representation of expansion segments ES27 (A) and ES7 (B) in *C. auris*, *C. albicans* and *S. cerevisiae*.

(B) The shoulder of the ES7 segment is indicated by a blue shadow with numbering of the starting nucleotide and ending nucleotide: *C. auris* G539-U557; *C. albicans* G628-U652; *S. cerevisiae* G589-U612.

The color indicates a *C. auris* nucleotide that differs from the *C. albicans* and *S. cerevisiae* nucleotide: green - altered nucleotide; blue - additional nucleotide.

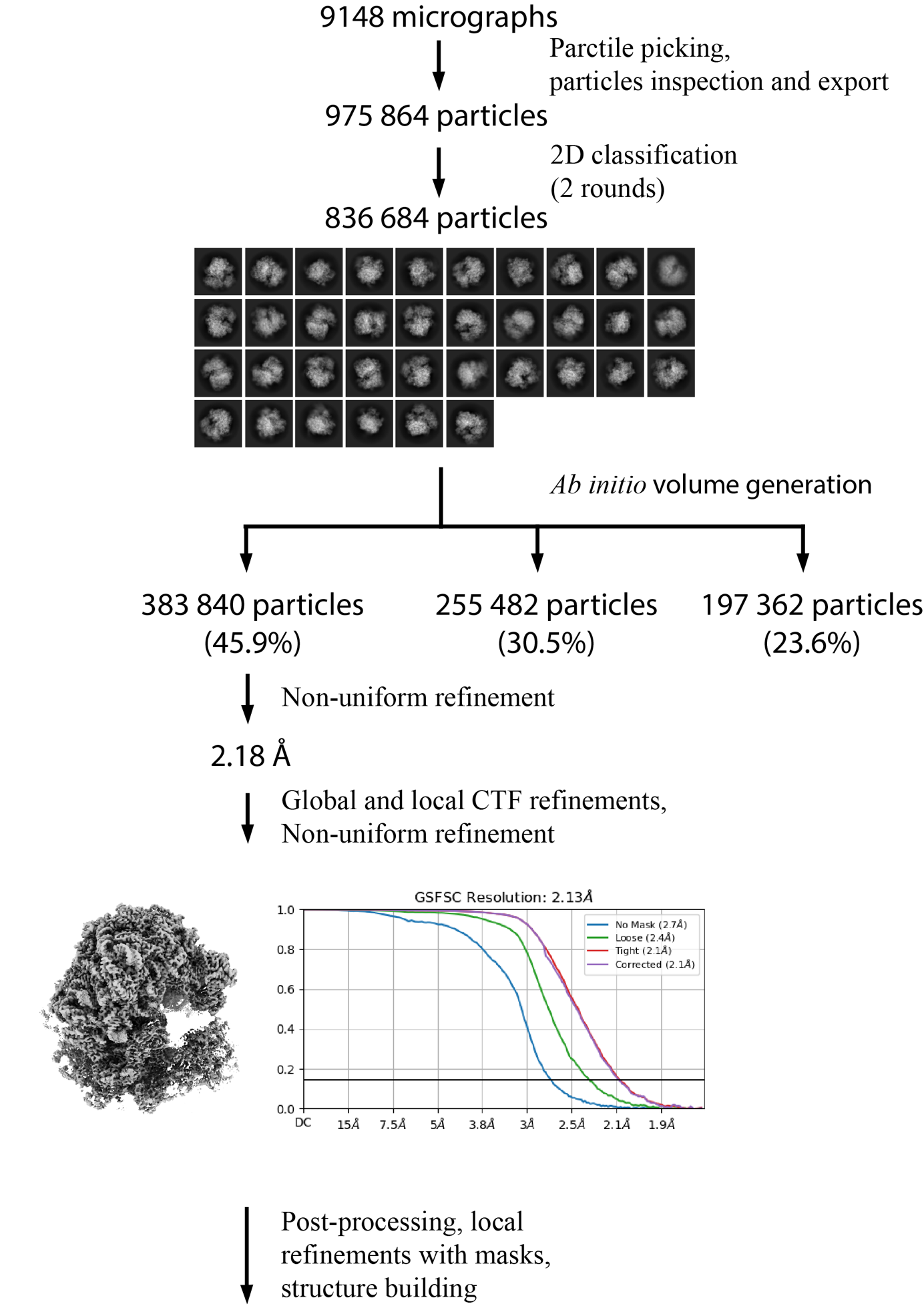

Fig. S5. Cryo-EM data processing scheme for the vacant *C. auris* ribosome.

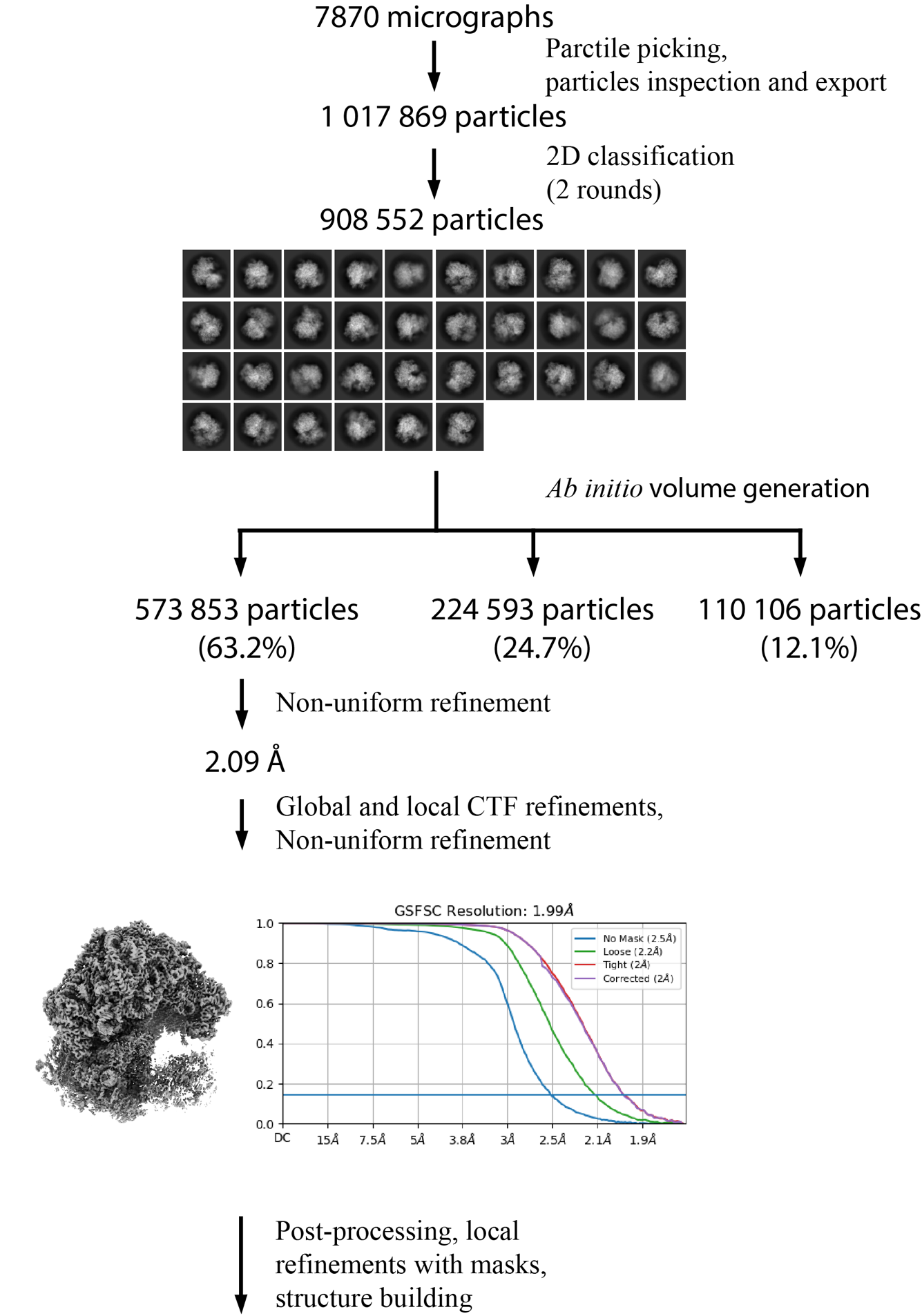

Fig. S6. Cryo-EM data processing scheme for the *C. auris* ribosome in complex with cycloheximide and geneticin.

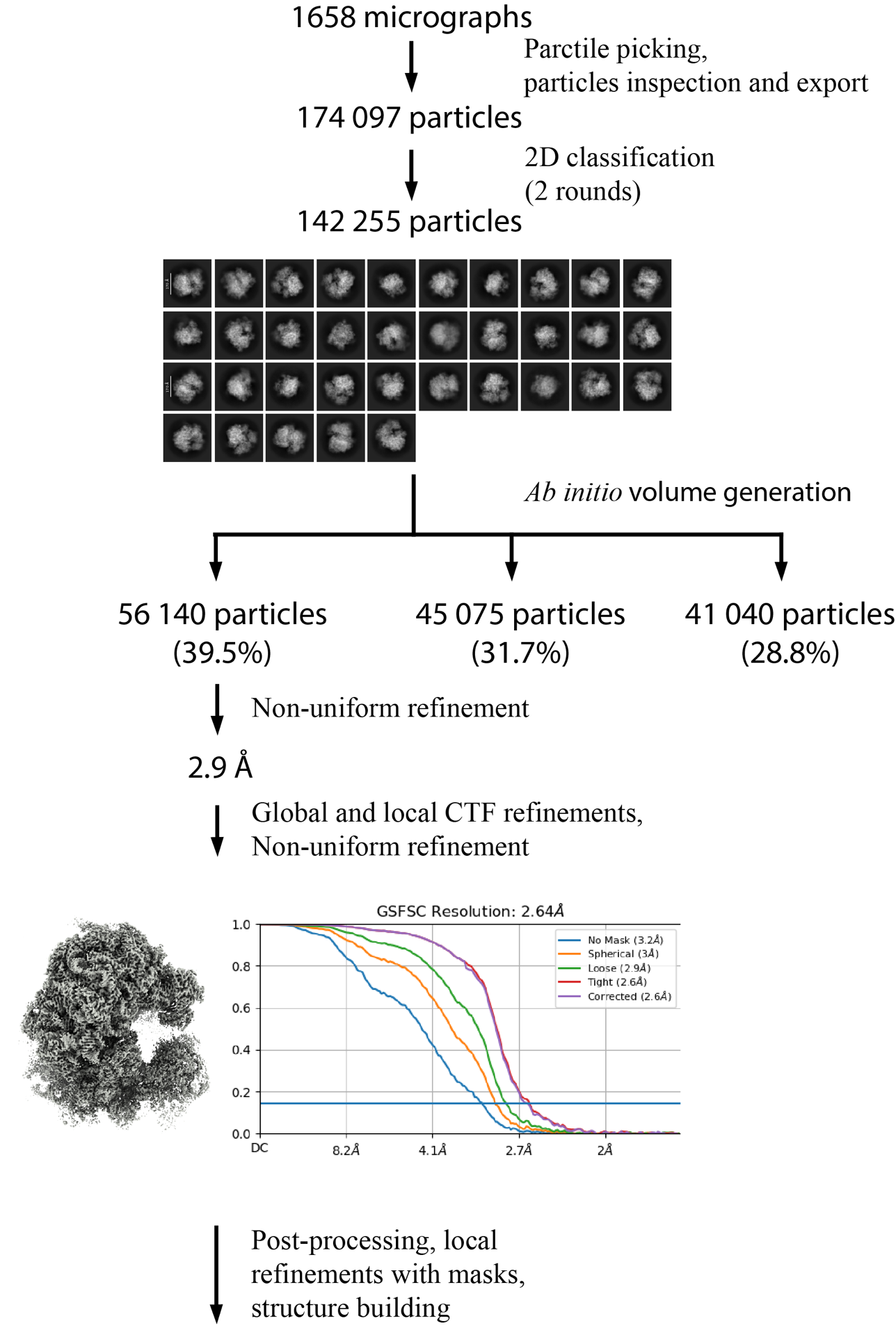

Fig. S7. Cryo-EM data processing scheme for the *C. auris* ribosome in complex with blasticidin S.

**References**

1. Toribio R, Díaz-López ,Irene, and Ventoso I. New insights into the topology of the scanning ribosome during translation initiation: Lessons from viruses. *RNA Biol* 2016;**13**:1223–7.

2. Díaz-López I, Toribio R, Berlanga JJ *et al.* An mRNA-binding channel in the ES6S region of the translation 48S-PIC promotes RNA unwinding and scanning. Sonenberg N, Manley JL, Sonenberg N (eds.). *eLife* 2019;**8**:e48246.

3. Anger AM, Armache J-P, Berninghausen O *et al.* Structures of the human and Drosophila 80S ribosome. *Nature* 2013;**497**:80–5.

4. Alkemar G, Nygård O. Secondary structure of two regions in expansion segments ES3 and ES6 with the potential of forming a tertiary interaction in eukaryotic 40S ribosomal subunits. *RNA* 2004;**10**:403–11.

5. Hariharan N, Ghosh S, Palakodeti D. The story of rRNA expansion segments: Finding functionality amidst diversity. *WIREs RNA* 2023;**14**:e1732.

6. Gulay SP, Bista S, Varshney A *et al.* Tracking fluctuation hotspots on the yeast ribosome through the elongation cycle. *Nucleic Acids Res* 2017;**45**:4958–71.

7. Fujii K, Susanto TT, Saurabh S *et al.* Decoding the Function of Expansion Segments in Ribosomes. *Mol Cell* 2018;**72**:1013-1020.e6.

8. Kišonaitė M, Wild K, Lapouge K *et al.* High-resolution structures of a thermophilic eukaryotic 80S ribosome reveal atomistic details of translocation. *Nat Commun* 2022;**13**:476.

9. Shedlovskiy D, Zinskie JA, Gardner E *et al.* Endonucleolytic cleavage in the expansion segment 7 of 25S rRNA is an early marker of low-level oxidative stress in yeast. *Ournal Biol Chem* 2017;**292**:18469–85.

10. Gómez Ramos LM, Degtyareva NN, Kovacs NA *et al.* Eukaryotic Ribosomal Expansion Segments as Antimicrobial Targets. *Biochemistry* 2017;**56**:5288–99.
