## Supplemental Table 1 for "Unveiling the Molecular Architecture of *Candida auris* Ribosome"

| **Atom number/Chain** | **Nucleotide/Chain/rRNA - Bond Length** | **Amino Acid/Protein - Bond Length (name of protein)** | **Other interactions** | **Location in the structure** |
| --- | --- | --- | --- | --- |
| **Large subunit** | | | | |
| 3289/A | 358A/A/25S – 3.08, 3.09, 358  360A/A/25S – 2.52, 3.16, 2.66, 3.04  393A/A/25S – 3.16  394C/A/25S – 2.65  395G/A/25S – 2.76, 2.69, 3.05  396U/A/25S – 2.86  397G/A/25S – 2.49 |  | Interaction with H2O | Surface of the large subunit |
| 3290/A | 212U/A/25S – 3.06  359G/A/25S – 3.03  21C/C/5.8S – 2.83  22U/C/5.8S – 3.2 |  | Interaction with H2O | Surface of the large subunit |
| 3291/A | 421G/A/25S – 2.83, 2.89  422C/A/25S – 2.98  570U/A/25S – 2.25  571G/A/25S – 2.93  572G/A/25S – 2.86 | 60ARG/h – 3.06 (L33) |  | Surface of the large subunit |
| 3292/A | 450C/A/25S – 2.58  452U/A/25S – 2.79  453C/A/25S – 2.92  536C/A/25S – 2.62 | 70LYS/h – 2.47 (L33) | Interaction with H2O | Surface of the large subunit |
| 3293/A | 479G/A/25S – 3.03, 3.16  480G/A/25S – 3.07, 3.18  481G/A/25S – 3.11, 2.25, 2.99  482A/A/25S – 2.64  505U/A/25S – 2.83  506U/A/25S – 3.18, 3.07  510G/A/25S – 2.79 |  |  | Surface of the large subunit |
| 3294/A | 464G/A/25S – 3.04, 3.14  465G/A/25S – 2.87  466G/A/25S – 3.03  523G/A/25S - 3  524A/A/25S – 3.06 |  | Interaction with H2O | Surface of the large subunit |
| 3295/A | 1305U/A/25S - 3.06  1286C/A/25S – 2.54  1287C/A/25S – 2.84  1288G/A/25S – 3.09, 3.12 |  | Interaction with H2O | Surface of the large subunit |
| 3296/A | 563G/A/25S – 3.09  564G/A/25S – 2.51, 2.79, 2.86  565C/A/25S – 2.91  445G/A/25S – 2.76, 2.91  444G/A/25S – 3.12, 2.91 |  |  | Surface of the large subunit |
| 3297/A | 665G/A/25S – 2.87, 2.52  666G/A/25S – 2.71  686G/A/25S – 2.32, 3.17  687C/A/25S – 2.63  692U/A/25S – 3.17 | 73GLN/S – 2.8 (L10) | Interaction with H2O | Surface of the large subunit |
| 3298/A | 754A/A/25S – 3.03  851G/A/25S – 2.93  852G/A/25S – 2.8, 3.13, 3.01  870A/A/25S – 2.87, 2.6  871C/A/25S – 2.81, 3.16  872C/A/25S – 3.14  2330G/A/25S – 2.82, 3.11, 3.07  2510A/A/25S – 2.92  2511G/A/25S – 2.88, 2.8 |  | Interaction with H2O | Deep in the large subunit |
| 3299/A | 766A/A/25S – 3.03, 2.78  767C/A/25S – 2.78  769U/A/25S - 3  770G/A/25S – 3.15, 3.12, 2.74  771A/A/25S – 3.08  843U/A/25S – 3.16  844G/A/25S – 3.1, 3.13  845G/A/25S – 2.94 |  | Interaction with H2O | Deep in the large subunit |
| 3300/A | 779G/A/25S – 2.89  1864A/A/25S – 3.1  1872G/A/25S – 2.89, 2.92, 2.84, 3.16  1874U/A/25S – 3.11 |  | Interaction with H2O | Deep in the large subunit |
| 3301/A | 2053U/A/25S – 3.19, 2.86  2056U/A/25S – 2.59  2711C/A/25S – 3.15  2712A/A/25S – 2.8 |  | Interaction with H2O | Deep in the large subunit |
| 3302/A | 812C/A/25S – 3.12  813G/A/25S – 2.51  814G/A/25S – 2.73, 3.13, 3, 3.11, 3.03, 3.06  830C/A/25S – 2.71, 3.12  831G/A/25S – 2.91, 3.17, 2.97 |  | Interaction with H2O | Deep in the large subunit |
| 3303/A | 931A/A/25S – 3.16  927A/A/25S – 3.05, 2.46  1011G/A/25S – 2.82, 3.16  1045G/A/25S – 2.64  1048U/A/25S – 2.74, 3.02 |  | Interaction with H2O | Surface of the large subunit |
| 3304/A | 1012A/A/25S – 3.01, 3.08  1013A/A/25S – 2.83  1044C/A/25S – 2.47, 2.86, 2.73  1045G/A/25S – 2.67 |  | Interaction with H2O | Surface of the large subunit |
| 3305/A | 1022U/A/25S – 3.2, 2.95  1025U/A/25S – 8.06  1026U/A/25S – 3.17  1029U/A/25S – 3.17  1030U/A/25S – 2.78, 2.85  1031G/A/25S – 2.88 |  |  | Surface of the large subunit |
| 3306/A | 1140C/A/25S – 3.1, 3, 3.15  1151A/A/25S – 3.18  1242G/A/25S – 2.95, 3.1  1245C/A/25S – 2.05  1246C/A/25S – 2.69, 3.2  1248G/A/25S – 2.85, 2.86  1249A/A/25S – 3.05  91A/B/5S – 3.19, 2.33, 3.18, 3.15 |  | Interaction with H2O | Surface of the large subunit??? |
| 3307/A | 1290C/A/25S – 2.35  1300G/A/25S – 2.71  1301C/A/25S – 2.76, 3.09, 2.69 |  | Interaction with H2O | Surface of the large subunit |
| 3308/A | 1321C/A/25S – 2.7  1322G/A/25S – 3.03, 3.19  1323U/A/25S – 2.5, 2.86  1368C/A/25S – 3.09 |  | Interaction with H2O | Surface of the large subunit |
| 3309/A | 1386U/A/25S – 3.06  1387G/A/25S – 2.47, 2.81, 2.7  1389U/A/25S – 2.87  2273A/A/25S – 2.49, 3.03 |  | Interaction with H2O | Deep in the large subunit |
| 3310/A | 1429G/A/25S – 2.8  1458G/A/25S – 2.87, 3.18, 2.94, 3.06  1781G/A/25S – 2.64  1784A/A/25S – 3.03 |  | Interaction with H2O | Deep in the large subunit |
| 3311/A | 1476C/A/25S – 3.15  1477U/A/25S – 2.38  1480C/A/25S – 3.02  1526G/A/25S – 2.56, 3.05, 2.34  1527G/A/25S – 3.19, 2.96  1528G/A/25S – 3.04, 2.73 |  | Interaction with H2O | Deep in the large subunit |
| 3312/A | 1804G/A/25S – 2.46  1805U/A/25S – 3.15, 3.1 | 80ARG/T – 3.11, 3.04, 2.98 (L19)  83ALA/T – 2.96, 2.95, 2.87 (L19)  84ARG/T – 2.92, 3.13 (L19)  87ALA/T – 3.1 (L19) | Interaction with H2O | Surface of the large subunit |
| 3313/A | 2335A/A/25S – 3.17, 2.9, 3  2336C/A/25S – 2.99, 2.81, 3.11  2504G/A/25S – 3.1  2505U/A/25S – 2.99  2506G/A/25S – 3.1, 2.98, 2.57 |  | Interaction with H2O | Surface of the large subunit |
| 3314/A | 2566U/A/25S – 3. 2.87  2587A/A/25S – 2.28  2588G/A/25S – 3.02, 2.93  2590G/A/25S – 2.58  2603A/A/25S – 3.19, 2.51, 2.92  2604A/A/25S – 3.16 |  | Interaction with H2O | Surface of the large subunit |
| 3315/A | 1249A/A/25S – 3.03, 2.76  2788A/A/25S – 2.99, 3.05  2789U/A/25S – 3.1  2808G/A/25S – 2.52  2809G/A/25S – 2.64 |  |  | Surface of the large subunit |
| 3316/A | 2817U/A/25S – 2.86  2818G/A/25S – 2.89  2819G/A/25S – 3.1, 3.18, 3.14, 2.46  2827A/A/25S – 3.18  2836U/A/25S – 2.76  2837A/A/25S – 2.54, 3.1  2838G/A/25S – 2.59 |  | Interaction with H2O | Surface of the large subunit |
| 3317/A | 2898C/A/25S – 3.2  2901U/A/25S – 2.75  3047G/A/25S – 2.83  3048A/A/25S – 2.92  3049C/A/25S – 2.77 |  | Interaction with H2O | Surface of the large subunit |
| 3318/A | 3003G/A/25S – 2.92  3004G/A/25S – 2.81  3031A/A/25S – 2.63, 2.9 |  | Interaction with H2O | Surface of the large subunit |
| 3319/A | 3212A/A/25S – 2.59  3211A/A/25S – 3.17, 2.98  3265A/A/25S – 2.84  3269U/A/25S – 3.17 | 169THR/E – 3.07, 3.17 (L3)  170PRO/E – 2.75, 3.19 (L3)  171LEU/E – 3.07 (L3) | Interaction with H2O | Surface of the large subunit |
| 3320/A | 1417G/A/25S – 2.71  1418A/A/25S – 2.72  1419A/A/25S – 3.04  1420G/A/25S – 2.84, 2.72  1820U/A/25S – 3.09, 2.84 |  | Interaction with H2O | Surface of the large subunit |
| 201/B | 66G/B/5S – 2.46, 2.44, 2.96  67G/B/5S – 3.19, 3.01, 2.51  68G/B/5S – 3.15, 2.96, 3.19 |  |  | Surface of the large subunit |
| 201/C | 64U/C/5.8S – 2.65  67U/C/5.8S – 2.89  84A/C/5.8S – 2.94, 3.14, 2.64  85A/C/5.8S – 2.85, 3.19  89U/C/5.8S – 2.95 |  | Interaction with H2O | Surface of the large subunit |
| 202/C | 49G/C/5.8S – 3.17, 2.6, 2.85  51G/C/5.8S – 2.34, 3.16  62C/C/5.8S – 2.98  63G/C/5.8S – 3.14, 2.49  86U/C/5.8S – 2.53, 3.06 |  |  | Surface of the large subunit |
| 203/C | 74U/C/5.8S – 3.04, 3.13  76C/C/5.8S – 2.81  77A/C/5.8S – 2.79, 3.15  78G/C/5.8S – 3.06 | 81GLN/A/25S – 3.03 (L26) | Interaction with H2O | Surface of the large subunit |
| 205/C | 5U/A/25S – 2.51  144G/C/5.8S – 2.44, 3.14  146G/C/5.8S – 2.82  148G/C/5.8S – 3.14 |  | Interaction with H2O | Surface of the large subunit |
| 8023/HO | 135U/C/5.8S – 3.06  22G/A/25S – 3.16  1484C/A/25S – 2.87  1487G/A/25S – 2.6  1488U/A/25S – 2.77  1489U/A/25S – 2.82  1492A/A/25S – 2.58 |  | Interaction with H2O | Deep in the large subunit |
| **Small subunit** | | | | |
| 1801/CA/18S | 1616C/CA/18S – 3.16  1617G/CA/18S – 3.08, 2.85, 2.2, 2.63  1618U/CA/18S – 3.04, 3.05  1619C/CA/18S – 2.87  1620G/CA/18S – 3.03  1729A/CA/18S – 3.09, 3.09  1731A/CA/18S – 3.13, 2.75  1733U/CA/18S – 3.17, 2.87 |  |  | Surface of the small subunit |
| 1802/CA/18S | 34G/CA/18S – 3.18  35U/CA/18S – 3.17  36C/CA/18S – 2.7, 3.17, 2.98  454G/CA/18S – 2.08, 3.01  456G/CA/18S – 3.15, 2.91, 3.15, 2.72 |  | Interaction with H2O | Surface of the small subunit |
| 1803/CA/18S | 22A/CA/18S – 2.35  23G/CA/18S – 2.79  459A/CA/18S – 2.8  585C/CA/18S – 3.14  586G/CA/18S – 3.19 |  | Interaction with H2O | Deep in the small subunit |
| 1804/CA/18S | 200U/CA/18S – 2.98  202U/CA/18S – 3.15, 2.66, 3.19  203A/CA/18S – 3.09  205U/CA/18S – 2.76  244U/CA/18S – 3.03 |  |  | Surface of the small subunit |
| 1805/CA/18S | 467A/CA/18S – 3.14, 2.37, 3.14, 2.95  468C/CA/18S – 3.14  495A/CA/18S – 2.98  530A/CA/18S – 2.5 |  | Interaction with H2O | Surface of the small subunit |
| 1806/CA/18S | 1051A/CA/18S – 2.35  1053G/CA/18S – 3.19  1055G/CA/18S – 3.12  1057A/CA/18S – 2.94, 2.4  1267G/CA/18S – 3.18  1268U/CA/18S – 2.56, 3.05, 3.19 | 156LYS/u (S2) – 3.13  158GLY/u – 3.05 (S2) | Interaction with H2O | Deep in the small subunit |
| 1807/CA/18S | 144U/CA/18S – 2.71, 3.18, 2.44  160A/CA/18S – 3.13  161G/CA/18S – 2.9  162A/CA/18S – 3.1 |  | Interaction with H2O | Surface of the small subunit |
| 1808/CA/18S | 395U/CA/18S – 2.65  410A/CA/18S – 2.94 |  | Interaction with H2O | Surface of the small subunit |
| 1809/CA/18S | 100U/CA/18S – 3.08  1379G/CA/18S – 2.78, 2.42, 3.11  1638G/CA/18S – 3.18, 2.28  1639C/CA/18S – 3.09  1708C/CA/18S – 3.16 | 27GLN/ZA – 3.1, 2.76, 3.12 (S8) | Interaction with H2O | Surface of the small subunit |
| 1810/CA/18S | 105U/CA/18S – 3.13  111A/CA/18S – 3.11  290A/CA/18S – 3.1  292U/CA/18S – 2.66  373G/CA/18S – 2.74 |  | Interaction with H2O | Deep in the small subunit |
| 1811/CA/18S | 1252U/CA/18S – 3.19  1253U/CA/18S – 2.94  1254C/CA/18S – 3.17  1256U/CA/18S – 3.18, 3.12  1397A/CA/18S – 2.68, 3.06  1398A/CA/18S – 3.16, 2.3  1399U/CA/18S – 3.18 |  |  | Deep in the small subunit |
| 74/OX | 39A/CA/18S – 2.72  40A/CA/18S – 2.93  41A/CA/18S – 3.19, 2.46, 2.99  427A/CA/18S – 2.76  454G/CA/18S – 2.72, 2.82  456G/CA/18S – 3.02 |  | Interaction with H2O | Surface of the small subunit |
| 75/OX | 59C/CA/18S – 2.72  60U/CA/18S – 2.7 |  |  | Surface of the small subunit |
| 76/OX | 100U/CA/18S – 2.99, 2.43, 2.93  346G/CA/18S – 3.09, 2.31  347U/CA/18S – 2.67  348A/CA/18S – 3.17, 3.07, 3.14  1637G/CA/18S – 3.16 |  | Interaction with H2O | Surface of the small subunit |
| 77/OX | 847G/CA/18S – 3.01  848G/CA/18S – 3.09  917U/CA/18S – 2.81, 3.05 |  | Interaction with H2O | Surface of the small subunit |
| 78/OX | 1087U/CA/18S – 3.17, 2.91  1088G/CA/18S – 2.86  1089G/CA/18S – 3.03, 2.59  1097G/CA/18S – 2.8, 3.18  1099U/CA/18S – 3.03, 2.71 |  | Interaction with H2O | Deep in the small subunit |
| 79/OX | 1569G/CA/18S – 3.19  1571C/CA/18S – 2.52, 2.65  1574C/CA/18S – 2.88 |  |  | Surface of the small subunit |
| 80/OX | 1646A/CA/18S – 2.96  1647G/CA/18S – 2.68  1701G/CA/18S – 3.19  1702G/CA/18S – 2.79  1705A/CA/18S – 3.19, 3.14, 2.57, 3.09 |  | Interaction with H2O | Surface of the small subunit |
| 81/OX | 619A/CA/18S – 2.95, 3.17, 3.05, 2.83  621U/CA/18S – 2.47  622U/CA/18S – 3.19  936A/CA/18S – 3.16 | 127ARG/ZE – 2.68 (S13) | Interaction with H2O | Surface of the small subunit |
