## Supplemental rRNA sequence for "Unveiling the Molecular Architecture of *Candida auris* Ribosome"

**LSU:**

TCGCCTCAAATCAGGTAGGACTACCCGCTGAACTTAAGCATATCAATAAGCGGAGGAAAAGAAACCAACAGGGATTGCCTCAGTAACGGCGAGTGAAGCGGCAAGAGCTCAACTTTGGAATCGCTCCGGCGAGTTGTAGTCTGGAGGTGGCCACCACGAGGTGTTCTAGCAGCAGGCAAGTCCTTTGGAACAAGGCGCCAGCGAGGGTGACAGCCCCGTACCTGCTTTTGCTAGTGCTTCCTGTGGCCCACCGACGAGTCGAGTTGTTTGGGAATGCAGCTCTAAGTGGGTGGTAAATTCCATCTAAGGCTAAATATTGGCGAGAGACCGATAGCGAACAAGTACAGTGATGGAAAGATGAAAAGCACTTTGAAAAGAGAGTGAAACAGTACGTGAAATTGTTGAAAGGGAAGGGCTTGCACCCAGACACGGTTTCGGCCGGGCCAGCATCAAGTAGAACGGGGTTAAAAGACCTGGGGAATGTAGCTACcTCttGGTAGTGTTATAGCCCTTgGGTGATGACCCCTGTTTTGCTTGAGGACAGCGGTCTCTAGGATGCTGGCGCAATGGTTGCAAGCCACCCGTCTTGGAACACGGACCAAGGAGTCTAACGTCTATGCGAGTGTACGGGTGAAAAACCCCTGCGCGGAATGAAAGTAAGAGGTTGGAACCCTCTCGGGGGTGCACAATCGACCGACCCAGAAGTGCTCGGACGGGTTTGAGTAGGAGCATAGCTGTTGGGACCCGAAAGATGGTGAACTATACCTGAATAGGGTGAAGCCAGAGGAAACTCTGGTGGAGGCTCGTAGCGGTTCTGACGTGCAAATCGATCGTCGAATTTGGGTATAGGGGCGAAAGACTAATCGAACCATCTAGTAGCTGGTTCCTGCCGAAGTTTCCCTCAGGATAGCAGAAGCTCGTTACAAACAGTTTTATGAGGTAAAGCGAATGATTAGAGGTCTCGGGGCTGAAATGGCCTTAGCCTATTCTCAAACTTTAAATATGTAAGAAGTCCTTGTTGCTTAATTGAACGTGGACATACGAATGTAGAGCTTTTAGTGGGCCATTTTTGGTAAGCAGAACTGGCGATGCGGGATGAACCGAACGCGAAGTTAAAGTGCCGGAATGCACGCTCATCAGACACCACAAAAGGTGTTAGTTCATCTAGACAGCCGGACGGTGGCCATGGAAGTCGGAATCCGCTAAGGAGTGTGTAACAACTCACCGGCCGAATGAACTAGCCCTGAAAATGGATGGCGCTCAAGCGTGCTACTTATACTTCGCCGGCAGCCTGTTTGCAGGTTGCCGAGTAGGCGGGCGTGGGGGTGGTGACGAAGCCTTGGCTGTGAAGCTGGGTCGAACCGCCCCTAGTGCAGATCTTGGTGGTAGTAGCAAATATTCAAACGGGAACTTTGAAGACTGAAGTGGGGAAAGGTTCCATGTCAACAGCAGATGGACATGGGTGAGTCGATCCTAACCCTCAGGTTAAACCTGATGAAAGTCGCCTCTTTGTTGAGGCCTCTGGGGAAAGGGAATCTGGTTAAAATTCCAGAACTTGGATGCGGAACTCACGGCAACGTAACTGAATGTGGAGACGCCGGCGTGGGCCCTGGGAGGAGTTTTCTTTTTCTTCTAACAGCCTGTGACCCTGGAATTGGATTATCCGGAGATGGGGTTTGTTGGCTGGAAGAGCGCGGCCTTTGTTTTGCCGCGTCtaGTGCGCCTACGACGGTCCTTGAAAATCCGCAGGAAGGAATGTTTTCGCGCCAAGTCGTACTGATAACCGCAGCAGGTCTCCAAGGTTAACAGCCTCTAGTTGATAGAACAATGTAGATAAGGGAAGTCGGCAAAATGGATCCGTAACTTCGGGATAAGGATTGGCTCTAAGGGTTGGGTGGTTTAGTGTTTCTTGCGGCAGCGGTGGTTGCCGGCCCAGGCGGCTAACCTTTCGGGGTCGGCTGCTTGGGCTGGCAACTCCGCCAAAGCTTGGAGCGCCGCCACCATTTGCAACCAACTTAGAACTGGTACGGACAAGGGGAATCTGACTGTCTAAtTTAAACATAGCATTGCGATGGTCAGAAAGTGATGTTGACGCAATGTGATTTCTGCCCAGTGCTCTGAATGTCAAAGTGAAGAAATTCAACCAAGCGCGGGTAAACGGCGGGAGTAACTATGACTCTCTTAAGGTAGCCAAATGCCTCGTCATCTAATTAGTGACGCGCATGAATGGATTAACGAGATTCCCACTGTCCCTATCTACTATCTAGCGAAACCACAGCCAAGGGAACGGGCTTGGCAGAATCAGCGGGGAAAGAAGACCCTGTTGAGCTTGACTCTAGTTTGACATTGTGAAAAGACATGGAGGGTGTAGAATAAGTGGGAGCTTCGGCGCCGGTGAAATACCACTACCTCCATTGTTTTTTTACTTACTGGGTGAAGGAGAGCTGGTCGCGAGACCAATTTCTTGCTTTGCATTTTTTCTCCGTCCAAGACATTGTCAGGTGGGGAGTTTGGCTGGGGCGGCACATCTGTaAAACGATAACGCAGGTGTCCTAAGGGGGGCTCATGGAGAACAGAAATCTCCAGTGGAACAAAAGGGTAAAAGCCCCCTTGATTTTGATTTTCAGTGTGAATACAAACCATGAAAGTGTGGCCTATCGATCCTTTAGTCCCTCGGAATTTGAGGCTAGAGGTGCCAGAAAAGTTACCACAGGGATAACTGGCTTGTGGCAGTCAAGCGTaCATAGCGACATTGCTTTTTGATTCTTCGATGTCGGCTCTTCCTATCATACCGAAGCAGAATTCGGTAAGCGTTGGATTGTTCACCCACTAATAGGGAACGTGAGCTGGGTTTAGACCGTCGTGAGACAGGTTAGTTTTACCCTACTGATGGACCGTTGTTGCAATAGTAATTGAACTTAGTACGAGAGGAACCGTTCATTCAGATAATTGGTATTTGGCCCTGTCTGACCAGGCACCGGGCCGAAGCTACCATCTGCTGGATTATGGCTGAACGCCTCTAAGTCAGAATCCATGCTAGACGCGACGACAATTTTTGGCCTCGCCTGCTGCTAGTTGGATACGAATAAGCTACGGCGCGGAACCATACAAGGTGGTGTTTGCTGGGTCACGGAAAGGTGGCCTGGTGGCCACGAATTGCAATGTCACACGGGCGGGGATGGATCCTTTGCATACGACTTAGATGTGCAACGGGGTATTGTAAGCGGTAGAGTAGCCTTGTTGTTACGATCCGCTGAGATTAAGCCTCTGTTGTCGGGTTTGT

**SSU:**

TATCTGGTTGATCCTGCCAGTAGTCATATGCTTGTCTCAAAGATTAAGCCATGCATGTCTAAGTATAAACGATTATACAGTGAAACTGCGAATGGCTCATTAAATCAGTTATCGTTTATTTGATAGTACCTTGCTAATTGGCATACGCCTGGCAATTCTAGAGCTAATACATGCGCAAAAGCCCGACTTCTGGGAGGGCTGTATTTATTAGATAAAAAATCAACGCTGCTTGATGATTCATAATAACTTGTCGAATCGCATGGCCTCGTGCCGGCGATGGTTCATTCAAATTTCTGCCCTATCAACTTTCGATGGTAGGATAGAGGCCTACCATGGTTTCAACGGGTAACGGGGAATAAGGGTTCGGTTCCGGAGAGGGAGCCTGAGAAACGGCTACCACATCCAAGGAAGGCAGCAGGCGCGCAAATTACCCAATCCCGACACGGGGAGGTAGTGACAATAAATAACGATGCAGGGCCCTTTCGGGTCTTGTAATTGGAATGAGTACAATGTAAATACCTTAACGAGGAACAATTGGAGGGCAAGTCTGGTGCCAGCAGCCGCGGTAATTCCAGCTCCAAGAGCGTATATTAAAGTTGTTGCAGTTAAAAAGCTCGTAGTTGAACCTTGGGTTTTGGAGGGAGGTCCACCTCACGGTGAGTACTTCCATATCCAAGACCTTTCCTCTGCTTCCTCGCAAGAGGCAGCAGAATATTACTTTGAGTAAATGAGAGTGTTCAAAGCAGGCGCACGCTTGAATCTGTTAGCATGGAATAATAGAATAGGACGCATGGTTCTATTTTGTTGGTTTCTAGGACCATCGTAATGATTAATAGGGACGGTCGGGGGCATCAGTATTCAGTTGTCAGAGGTGAAATTCTTGGATTTACTGAAGACTAACTACTGCGAAAGCATTTGCCAAGGACGTTTTCATTAATCAAGAACGAAAGTTAGGGGATCGAAGATGATCAGATACCGTCGTAGTCTTAACCATAAACTATGCCGACTAGGGATCGGGCGGCGTTCATCTAGTGACGCGCTCGGCACCTTACGAGAAATCAAAGTTTTTGGGTTCTGGGGGGAGTATGGTCGCAAGGCTGAAACTTAAAGGAATTGACGGAAGGGCACCACCAGGAGTGGAGCCTGCGGCTTAATTTGACcCAACACGGGGAAACTCACCAGGTCCAGACACAATAAGGATTGACAGATTGAGAGCTCTTTCTTGATTTTGTGGGTGGTGGTGCATGGCCGTTCTTAGTTGGTGGAGTGATTTGTCTGCTTAATTGCGATAACGAACGAGACCTTAACCTCTAAATAGTGCTGCCAGCGCGCCTCTTGTGCTGGCGCTGCACTTCTTAGGGGGACTATTGACTTGAAGTCGATGGAAGTTTGAGGCAATAACAGGTCTGTGATGCCCTTAGACGTTCTGGGCCGCACGCGCGCTACACTGACGGAGCCAGCGAGTTGACCTTGGCCGAGAGGCCTGGGAAATCTTGGGAAACTCCGTCGTGCTGGGGATAGAGCATTGCAATTGTTGCTCTTCAACGAGGAATTCCTAGTAAGCGCAAGTCACCAACTTGCGTTGATTACGTCCCTGCCCTTTGTACACACCGCCCGTCGCTACTACCGATTGAATGGCTTAGTGAGGCCTCCGGATCTGGCATGCCCCGAGGGCAACCTCGCCGCGCGCCGGAGAAGCTGGTCAAACTTGGTCATTTAGAGGAAGTAAAAGTCGTAACAAGGTTTCCGTAGGTGAACCTGCGGAAGGATCATTA
